# PUMILIOs and m^6^A-ECT2/ECT3 share mRNA targets and exert opposing control over organogenesis

**DOI:** 10.1101/2025.10.11.681827

**Authors:** Junyu Chen, Marina Sangés Ametllé, Sarah Rennie, Laura Arribas-Hernández, Peter Brodersen

## Abstract

PUMILIO/Fem-3 Binding Factor (PUF) proteins are conserved eukaryotic mRNA-binding proteins. Higher plants encode an expanded family of PUF proteins with 26 members in *Arabidopsis thaliana* of which 6 (PUM1-PUM6) have a domain organization equivalent to metazoan PUF proteins implicated in mRNA regulation. Here, we show that the PUF proteins PUM2 and PUM6 are expressed in mitotically active cells of root and shoot meristems, while PUM5 is mostly expressed in the differentiation zone of root meristems exiting from active division. Increased PUM2 dosage in proliferating cells causes growth repression and developmental delay reminiscent of mutants with defects in cytoplasmic YTHDF proteins that bind to *N6*-methyladenosine (m^6^A) in mRNA. Remarkably, even weak defects in the m^6^A-YTHDF system enhance the effect of increased PUM2 dosage. In line with this genetic interaction, mRNA targets overlap significantly between PUFs and the major YTHDF proteins ECT2 and ECT3. Furthermore, the enrichment of the core PUF recognition motif URUAY around m^6^A sites is largely explained by PUF targeting, not by direct URUAY methylation as previously suggested. Our results indicate that the balance between the reciprocal activities of the deeply conserved cytoplasmic RNA binding protein families PUF and YTHDF is an important element of growth control in plants.

## INTRODUCTION

RNA-binding proteins (RBPs) govern pre-mRNA processing steps and mature mRNA properties, including nuclear export, localization, translation and decay. Similar to transcription factor families such as MYB, homeobox, and NF-Y which play central roles in eukaryotic genetic control, several RBP families are deeply conserved across eukaryotic phylogenetic clades. Examples include the cytoplasmic poly(A)-binding protein (PABP) with broad functions in mRNA translation and stability (Mangus et al., 2003; Eliseeva et al., 2013; Passmore and Coller, 2022), the Musashi family (Fox et al., 2015) recently found to be required for post-embryonic plant development (Kairouani et al., 2023), and the YTHDF proteins that mediate many plant growth-related functions of the covalent mRNA modification *N6*-methyladenosine (m^6^A) by direct m^6^A-binding (Brodersen and Arribas-Hernández, 2024).

Pumilio (PUM)/Fem-3 Binding Factor (FBF) proteins (PUFs) constitute another class of deeply conserved pan-eukaryotic RBPs (Najdrová et al., 2020). These proteins consist of a long N-terminal intrinsically disordered region (IDR) followed by a highly conserved globular PUM homology domain (PUM-HD). PUF proteins bind sequence-specifically to 3’-untranslated regions (3’-UTRs) of mRNAs (Wickens et al., 2002), and the sequence specificity relies on the PUM-HD that contains 8 PUF repeats (Zamore et al., 1997), each responsible for recognition of a single nucleobase (Wang et al., 2002). In mammals, the resulting 8-nt consensus Pumilio recognition element (PRE) is UGUANAUA (H = A/G/C/U) (White et al., 2001), or a less stringent version URUAWAUW (R = G/A, W = A/U) determined by transcriptome-wide binding site analysis of human Pum2 in cultured cells (Hafner et al., 2010). Similar partially degenerate PUF consensus sites with a UGUA core have emerged from transcriptome-wide identification and sequence analysis of PUF-bound mRNAs in *Drosophila melanogaster* (Drosophila) (Gerber et al., 2006; Laver et al., 2015) and the yeast *Saccharomyces cerevisiae* (Gerber et al., 2004), as well as direct *in vivo* binding site analysis in *Caenorhabditis elegans* (Prasad et al., 2016). The binding site degeneracy despite the stringent 1 PUF repeat:1 nucleobase recognition in pure PUF-PRE complexes is explained by the fact that the PUF-RNA interaction is generally assisted by other RBPs *in vivo*, such as the Zn-finger protein Nanos (Asaoka-Taguchi et al., 1999; Sonoda and Wharton, 1999) that, unlike PUFs themselves, is restricted to metazoans (De Keuckelaere et al., 2018). Compared to Pum-PRE complexes, the topology of the PUF repeats is adjusted in ternary Pum-Nanos-PRE complexes such that nucleobase interactions with the 5’-part of the PRE are strengthened while other bases flip away from direct contacts with PUF repeats (Weidmann et al., 2016), thus enabling *in vivo* recognition of a wider variety of PREs (Arvola et al., 2017). Similar molecular principles - albeit with different effects on affinity and specificity - apply to the germline stem cells of *C. elegans* where FBF interaction with PREs in target is assisted by the nematode-specific Zn-finger protein Lateral Signaling Target-1 (LST-1) (Qiu et al., 2019).

In addition to the 8 PUF repeats, canonical PUF proteins harbor two conserved sequences flanking the PUF repeats that are implicated in RNA binding and protein-protein interactions (Zhang et al., 1997; Kraemer et al., 1999). Such canonical PUF proteins are pivotal for multiple fundamental cell functions in metazoans and yeast. In Drosophila, Pumilio - the sole and founding member of the PUF family - is required for formation of abdominal segments in the early embryo (Lehmann and Nüsslein-Volhard, 1987; Barker et al., 1992), primarily via translational repression and deadenylation of the mRNA encoding the key transcription factor *hunchback* (*hb*) (Murata and Wharton, 1995; Wreden et al., 1997; Wharton et al., 1998). Translational repression and accelerated decay of target mRNAs mediated by direct interactions with key factors controlling cap-dependent translation initiation and deadenylation-dependent mRNA decay also underlie many other important biological functions of PUF proteins (Olivas and Parker, 2000; Cho et al., 2006; Deng et al., 2008; Cao et al., 2010; Chritton and Wickens, 2011; Blewett and Goldstrohm, 2012; Van Etten et al., 2012; Weidmann and Goldstrohm, 2012; Weidmann et al., 2014; Arvola et al., 2019; Enwerem et al., 2021; Haugen et al., 2022; Haugen et al., 2024). In metazoans, these functions include stem cell maintenance and differentiation (Crittenden et al., 2002; Ariz et al., 2009; Racher and Hansen, 2012; Leeb et al., 2014; Lin et al., 2018; Uyhazi et al., 2020), germ cell development (Zhang et al., 1997; Forbes and Lehmann, 1998; Asaoka-Taguchi et al., 1999; Kraemer et al., 1999; Chen et al., 2012; Mak et al., 2018), and neuronal development and function (Menon et al., 2004; Muraro et al., 2008; Gennarino et al., 2015; Zhang et al., 2017; Gennarino et al., 2018; Zahr et al., 2018; Martínez et al., 2019), and in yeast, PUFs control mitochondrial activity via glucose-regulated translation of target mRNAs encoding mitochondrial proteins (Gerber et al., 2004; García-Rodríguez et al., 2007; Saint-Georges et al., 2008). Interestingly, both metazoan and yeast PUFs have also been reported to translationally activate target mRNAs in some cases (Deng et al., 2008; Kaye et al., 2009; Suh et al., 2009; Lee and Tu, 2015; Bohn et al., 2017; Naudin et al., 2017), indicating that PUF functions cannot simply be designated as repressive or activating. Elegant studies of yeast PUF proteins have shown the existence of a repressor-to-activator switch driven by extensive IDR phosphorylation (Deng et al., 2008; Lee and Tu, 2015), a molecular principle that may apply more broadly.

In plants, the PUF family has undergone extensive expansion, with 26 genes belonging to five phylogenetic groups encoding PUF-repeat-containing proteins in the flowering plant *Arabidopsis thaliana* (Arabidopsis) (Francischini and Quaggio, 2009). Six of these 26 proteins, PUM1-PUM6, belong to the canonical class and stand out from the other phylogenetic groups because they exhibit >50% amino acid identity to Drosophila Pumilio in the PUF repeats, and display specific binding activity to the UGUACAUA-element from the 3’-UTR of the *hb* mRNA in yeast three-hybrid assays (Francischini and Quaggio, 2009). Little is known about the biological functions of these canonical plant PUM proteins, however. PUM5 exerts an antiviral function against the (+)-strand RNA virus Cucumber Mosaic Virus by direct binding to viral RNA containing the core PUM binding site UGUAY, perhaps via translational repression of viral mRNA (Huh et al., 2013). Attempts at endogenous mRNA target identification have so far been limited to yeast three-hybrid screening that identified a handful of mRNAs as likely direct PUM2 targets (Francischini and Quaggio, 2009). Systematic gain-of-function approaches have not been reported, and loss-of-function studies are complicated by the potentially redundant functions of the six *PUM1-PUM6* genes. In addition, the metazoan facilitators of PUF-RNA interaction Nanos and LST-1 do not have clear plant homologues, precluding insights into plant PUF function through genetic analysis of such factors. Thus, at present, little is known about endogenous functions and mRNA targets of the highly conserved canonical PUF proteins in plants. In many plant species, the URUAY (R = A/G, Y = U/C) motif is enriched in the vicinity of m^6^A in 3’-untranslated regions (3’-UTRs), and its similarity to the core PRE was noted in one of the earliest reports on m^6^A maps in rice and Arabidopsis (Li et al., 2014). Subsequently, however, the idea that URUAY might be a plant-specific methylation motif caught fire, fueled in particular by the enrichment found in pieces of RNA crosslinked with formaldehyde to the major cytoplasmic m^6^A-binding protein in Arabidopsis, the YTHDF protein EVOLUTIONARILY CONSERVED C-TERMINAL REGION2 (ECT2) (Wei et al., 2018). However, formaldehyde creates both protein-protein and protein-RNA crosslinks, resulting in the detection of indirect protein-RNA interactions. When methodology appropriate for transcriptome-wide identification of direct mRNA targets and binding sites of ECT2 was employed, ECT2 was found to bind to m^6^A primarily in the DRACH (D=G/A/U, R=G/A, H=U/A/C) context, although still with significant enrichment of URUAY (R=G/A, Y=C/U) at and around crosslinking sites (Arribas-Hernández et al., 2021b). Thus, while little evidence supports URUAY as a *bona fide* plant-specific m^6^A motif, its enrichment in m^6^A-containing 3’-UTRs remains an intriguing observation that calls for a molecular explanation.

Here, we present a functional investigation of canonical Arabidopsis PUF proteins, including transcriptome-wide target mRNA identification and an assessment of functional links between PUFs and m^6^A-YTHDF. We show that PUM2/6 expression overlaps with ECT2/3 expression in mitotically active cells and that the overlap in PUM2/5/6 and ECT2/3 mRNA targets is significant such that the enrichment of URUAY around m^6^A sites in ECT2/3 mRNA targets is largely driven by PUF-targeting. Increased dosage of PUM2 causes growth arrest reminiscent of inactivation of the m^6^A-YTHDF axis, and *ECT2* and *PUM2* interact genetically in growth control. These results suggest that balancing of the deeply conserved PUF and YTHDF proteins, with opposing biological function, is part of the decision-making on whether primordial cells grow and divide or exit the cell cycle.

## RESULTS

### PUM2/PUM6 and ECT2 expression overlap in mitotically active cells

To test whether PUF and ECT2/3 function could be related *in vivo*, we first determined the expression patterns of three canonical PUF proteins, PUM2, PUM5 and PUM6, using fluorescent protein fusions expressed under the control of the endogenous promoter and terminator sequences. These analyses showed that expression of PUM2 and PUM6 overlaps with that of ECT2 in mitotically active regions developing leaves and root apical meristems (RAM) (Fig. 1A,B). PUM5 expression, on the other hand, was strongest in the elongation zone of the RAM where ECT2/3 expression is low (Fig. 1A,B). These expression patterns determined by fluorescent protein fusions were confirmed by inspection of published mRNA-seq data from dissected tissues (Mergner et al., 2020) and single root cells (Denyer et al., 2019) that showed co-expression of PUM2 and PUM6 in proliferating cells of the RAM (Supplementary Fig. S1). Hence, the expression of the canonical PUFs PUM2 and PUM6 overlaps sufficiently with ECT2 in mitotically active cells to warrant further comparative functional analyses.

**Figure 1.**
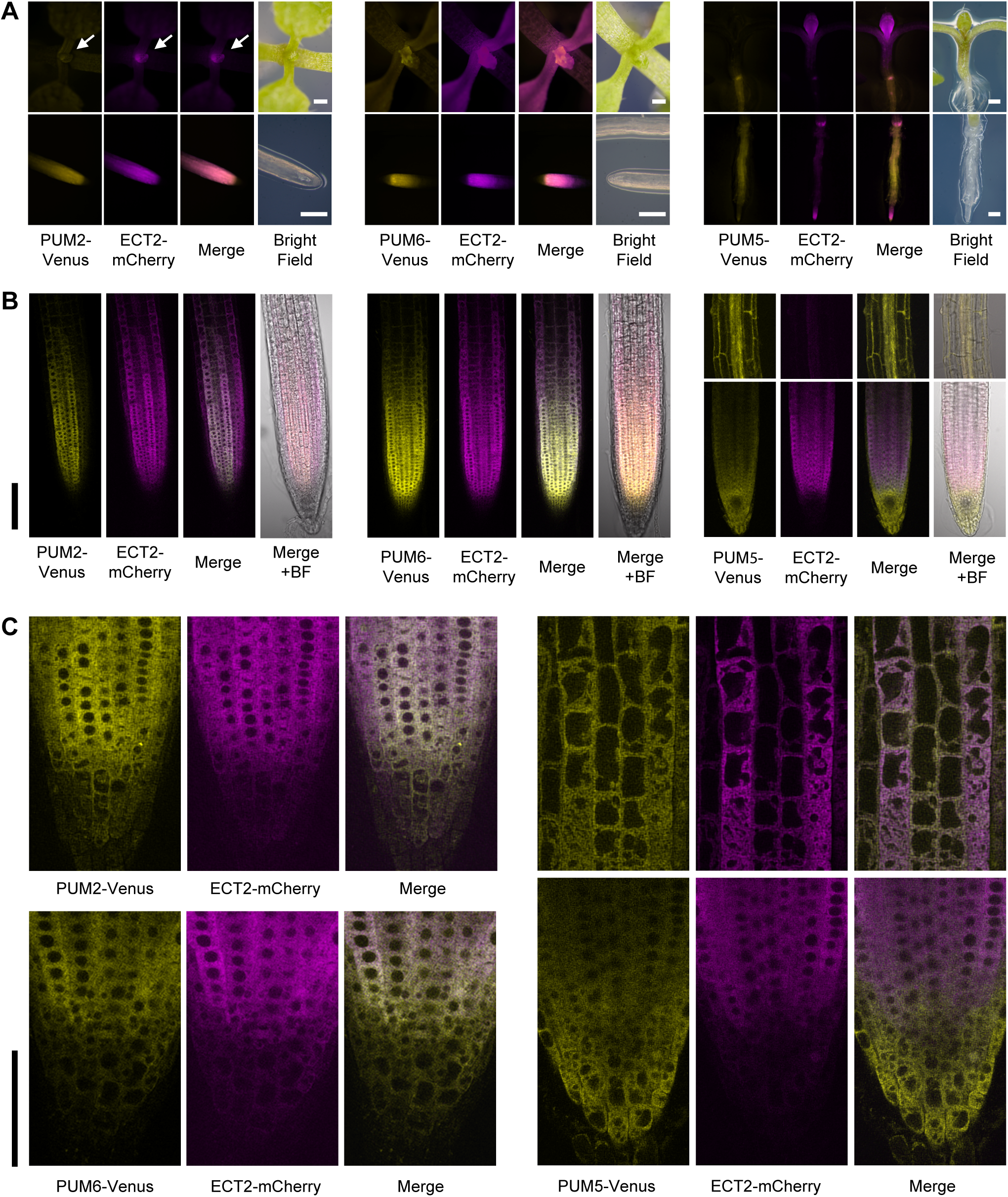
Expression patterns of PUM2, PUM5 and PUM6 compared to ECT2. **(A)** Fluorescence microscopy of aerial and root parts of 7-day-old Arabidopsis seedlings co-expressing PUM2/5/6-Venus and ECT2-mCherry. White arrows indicate the shoot apex. Scale bars, 100 μm. **(B-C)** Confocal fluorescence microscopy of seedling roots of the transgenic lines described in (A). Scale bars, 50 μm. BF, bright field.

### Canonical PUF proteins are largely cytoplasmic

We next used confocal microscopy to study the subcellular localization of the three fluorescently tagged PUF proteins in root cells. We observed fluorescence signal predominantly, if not exclusively, in the cytoplasm (Fig. 1C). Hence, as in other eukaryotic organisms (Macdonald, 1992; Zhang et al., 1997; Gerber et al., 2004; Morris et al., 2008), canonical PUF proteins in Arabidopsis are cytoplasmic.

### mRNA target identification of PUF proteins using HyperTRIBE

To acquire the next layer of basic functional information on PUF proteins, we sought to identify mRNA targets of PUM2, PUM5 and PUM6 transcriptome-wide. This information would also allow a judgment of mRNA target overlap with ECT2 and ECT3, and hence address more directly whether PUFs and ECTs could participate in related biological processes. We used the proximity labeling approach HyperTRIBE (HT), (McMahon et al., 2016; Rahman et al., 2018) in which a hyperactive variant of the catalytic domain of the Drosophila Adenosine Deaminase Acting on RNA (ADARcd^E488Q^, henceforth ‘ADAR’) is fused to the RBP of interest. mRNA sites with significantly higher adenosine-inosine (A-I) editing proportions, read as A-G mutations in mRNA-seq, than in plants expressing similar levels of FLAG-ADAR are then used to identify mRNA targets bound by the fusion protein. When searching for transgenic lines expressing PUM-FLAG-ADAR or FLAG-ADAR, we found that PUF expression levels were generally low. For PUM2 and PUM6, protein abundance was only high enough for immunoblot-based line selection in aerial tissues of seedlings, while only root tissues satisfied this criterion for PUM5 (Supplementary Fig. S2A). Hence, we used aerial tissue for target identification of PUM2 and PUM6, but roots for PUM5. Despite the use of endogenous promoter and transcription terminator sequences, *PUM2/5/6* transgenes were somewhat overexpressed compared to the endogenous levels in most of the selected lines with sufficient protein accumulation for immunoblot detection (Supplementary Fig. S2). We verified that this degree of overexpression did not produce growth phenotypes, nor did it appreciably alter the global gene expression profiles compared to the FLAG-ADAR-expressing controls (Supplementary Fig. S3).

We previously established procedures for target identification by HT of ECT2 and ECT3 in Arabidopsis (Arribas-Hernández et al., 2021b; Arribas-Hernández et al., 2021a). Using these procedures with five independent PUM2/5/6-FLAG-ADAR-and FLAG-ADAR-expressing lines (Supplementary Fig. S4), we found that principal component analysis based on editing proportions clearly separated samples prepared from PUM2/5/6-fused vs. free FLAG-ADAR lines (Supplementary Fig. S5A). Importantly, sites with significantly different editing proportions between the two genotypes (see Materials and methods) had higher editing proportions in PUM2/5/6-fused lines (Fig. 2A), as expected for *bona fide* target identification. These differentially edited sites defined 723 targets of PUM2, 2488 targets of PUM5, and 5273 targets of PUM6, with “targets” referring to distinct gene identifiers and hence not counting isoforms as distinct targets. Three additional properties support the conclusion that they are *bona fide* PUF mRNA targets. First, searches for 8-mer sites overrepresented in 3’-UTRs of PUM2/5/6 HT-targets clearly identified variants of the PUF consensus site (UGUAWHHW, W=A/U, H=U/A/C) as the most highly enriched 8-mer (Fig. 2B). Second, the distribution of PUM consensus binding sites showed clear bias towards 3’-UTRs in PUM2/5/6 target mRNAs compared to all expressed mRNAs (Fig. 2C). Third, overlaps between PUM2, PUM6 and PUM5 target sets were significant (p < 2.2×10^-16^ for chi-square test) despite the fact that different tissues were used for PUM5 and PUM2/6 (Fig. 2D). Indeed, PUM2 targets were nearly fully contained in the larger PUM6 target set (Fig. 2D), while most of the non-shared targets between PUM5 and PUM6 were explained by root-or shoot-specific expression (Fig. 2E). The significant overlaps in target sets are reassuring because of the high degree of sequence identity between the three PUM proteins. Finally, 5 of the 9 transcripts previously identified as PUM2 targets by yeast three-hybrid screening (Francischini and Quaggio, 2009) were also identified as HT-targets of at least one of the three PUF proteins studied here (Supplementary Fig. S5B).

**Figure 2.**
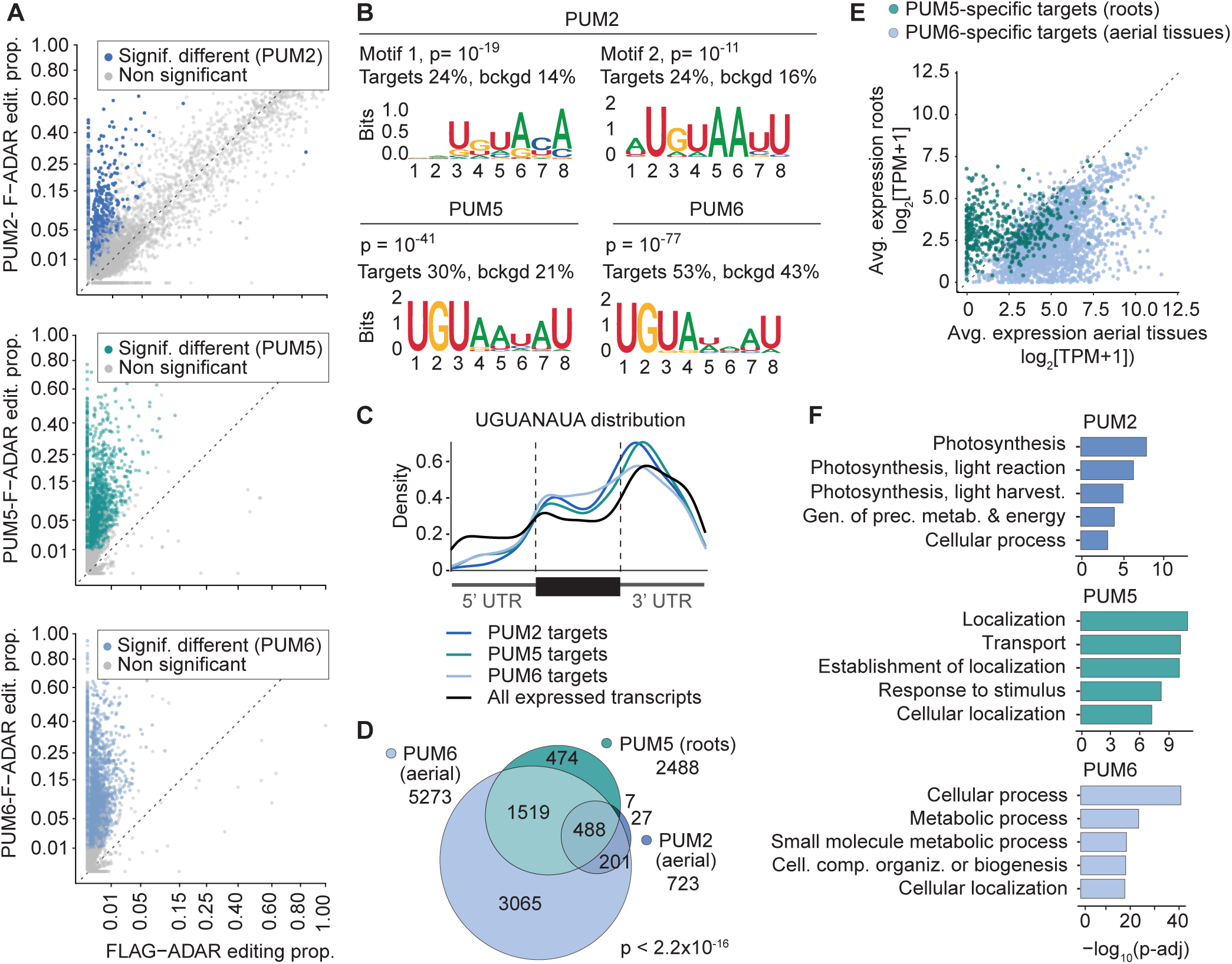
HyperTRIBE identification of PUM2/5/6 mRNA targets. **(A)** Scatter plots comparing the editing proportions in PUM2/5/6-FLAG-ADAR lines (y-axes) to those of the corresponding free FLAG-ADAR controls (x-axes). **(B**) 8-mer motifs enriched in the 3’-UTR of PUM2/5/6 targets obtained by HOMER-based *de novo* motif discovery. All 3’-UTR sequences in the Arabidopsis transcriptome were used as background for the algorithm. p reflects the significance of motif enrichment in PUM targets compared with background. ‘Target’ and ‘bkgrd’ stand for the percentages of sequence containing the given motif in PUM target group or background, respectively. **(C)** Metagene plot showing distribution of canonical PUM response elements (UGUANAUA) in PUM2/5/6 targets compared to all expressed transcripts as detected in PUM2/5/6 HyperTRIBE. **(D)** Overlap between PUM2, PUM5 and PUM6 target sets. The overlap between the three datasets was significantly greater than expected by chance (Chi-square test for mutual independence, p < 2.2×10^-16^). **(E)** Tissue-specific gene expression of PUM5-vs. PUM6-specific targets. **(F)** Gene Ontology (GO) enrichment analysis of PUM2/5/6 targets. All expressed genes in the corresponding HyperTRIBE samples were used as background.

Gene ontology analyses of the target sets showed that PUM2 targets are enriched in photosynthesis-related transcripts, while PUM6 targets include transcripts related to other metabolic processes (Fig. 2F). PUM5 targets contained metabolism-related transcripts shared with PUM6, but also showed a large set of targets enriched in stress response genes (categorized as ‘response to stimulus’) (Fig. 2F). We conclude that the HT approach identifies PUM2/5/6-bound transcripts with properties consistent with *bona fide* mRNA targets, and that prominent classes of PUM transcripts encode proteins involved in basic metabolic processes that include photosynthesis, nitrogen metabolism and oxidative phosphorylation.

We note that the difficulty of obtaining transgenic lines with perfectly balanced PUM6-FLAG-ADAR and FLAG-ADAR expression under the *PUM6* endogenous promoter (Supplementary Fig. S2) may cause more transcripts to be identified as PUM6 targets compared to PUM2 and PUM5. Hence, transcripts identified as PUF targets uniquely on the basis of the PUM6 HT experiment should be regarded as less high-confidence targets.

### PUM2/5/6 mRNA targets overlap substantially with those of m^6^A-ECT2/3

We next compared the PUM2/5/6 HT-targets to those previously identified for ECT2, ECT3 (Arribas-Hernández et al., 2021b; Arribas-Hernández et al., 2021a), their facilitators of m^6^A binding ALBA2 and ALBA4 (Reichel et al., 2024), as well as m^6^A targets identified by different single-nucleotide resolution techniques (Parker et al., 2020; Wang et al., 2024). In many cases, ECT2/3, ALBA2 and PUM2/5/6 HT-edited sites coincided on m^6^A-modified mRNAs (Fig. 3A). Indeed, we found that the overlaps with ECT2/3 mRNA targets were highly significant for all three PUF proteins in the relevant tissues (Chi-square test, p < 2.2×10^-16^ for all PUMs; Fig. 3B, Supplementary Fig. S6A,B). These overlaps were particularly pronounced for PUM2 and PUM6, whose expression patterns overlap tightly with those of ECT2 and ECT3 (Fig. 1A,B).

**Figure 3.**
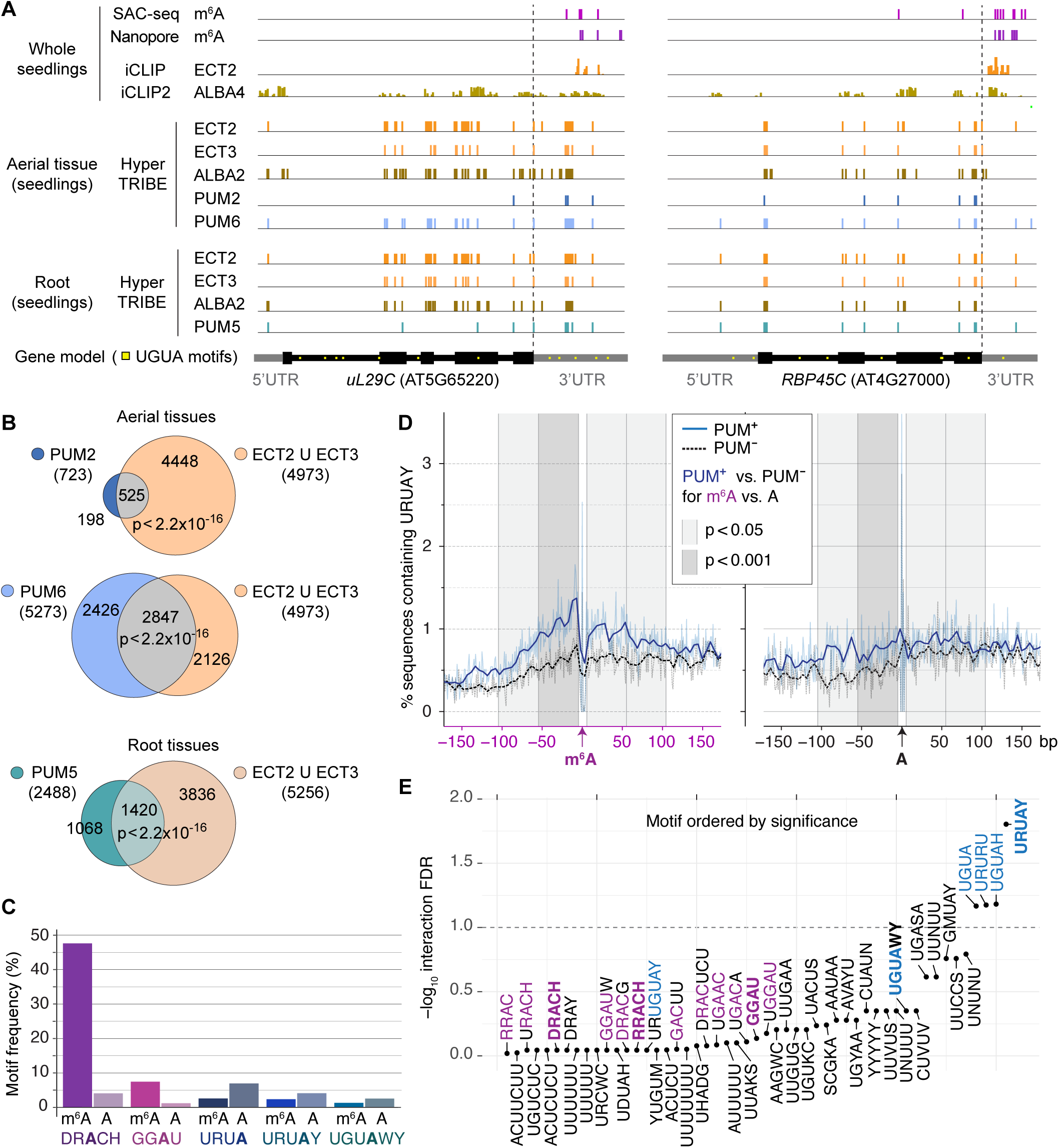
The overlap between PUM and ECT2/ECT3 target sets is substantial and drives the URUAY enrichment around m^6^A in ECT2/3 targets. **(A)** Examples of PUM HyperTRIBE (HT) targets, showing the location of significant PUM2/5/6 HT editing sites together with ECT2 iCLIP and ALBA4 iCLIP2 crosslink peaks (Arribas-Hernández et al., 2021b; Reichel et al., 2024), ECT2/3 HT editing sites (Arribas-Hernández et al., 2021b; Arribas-Hernández et al., 2021a), ALBA2 HT editing sites (Reichel et al., 2024), and m^6^A sites identified by Nanopore and m^6^A-SAC-seq (Parker et al., 2020; Wang et al., 2024). Potential PUM binding sites (UGUA) are also indicated on the gene model. **(B)** Overlap between PUM2/5/6 and ECT2/3 HT targets (Arribas-Hernández et al., 2021b; Arribas-Hernández et al., 2021a). The number of shared targets is significantly greater than expected by chance (Chi-square tests, p < 2.2×10^-16^). **(C)** Percentage of m^6^A sites in 3’-UTRs contained in various motifs previously found to be enriched around m^6^A (additional motifs are shown in Supplementary Fig. S6C). The m^6^A sites used for the analysis were determined at single-nucleotide resolution in seedlings by m^6^A-SAC-seq (Wang et al., 2024). Strong colors represent m^6^A sites and dim colors represent random As sampled in the 3’-UTRs of the same transcripts. The adenosine presumed to be methylated in each motif is highlighted in bold. **(D)** Percentage of 3’-UTR m^6^A-SAC-seq sites (Wang et al., 2024) (left panel) or random As in similar locations (right panel) in ECT2/3 target mRNAs with the URUAY (R=A/G, Y=C/U) motif present at given distances from the site, split by whether the transcript is PUM2/5/6 HT target (PUM^+^) or not (PUM^-^). Dark lines are smooth representations of the raw data (dim lines). Vertical bands in different shades of grey represent windows of 50-nt width around the site (excluding the immediately adjacent 5 nt). The associated p-values indicate the significance of the difference in URUAY depletion in the PUM^+^ to PUM^-^ subset comparisons with sequence windows centered either on verified m^6^A sites or random As. **(E)** Significance of the difference in motif depletion in the PUM^+^ to PUM^-^ subset comparisons using sequence windows centered on verified m^6^A sites and random As, given p-values corrected for multiple testing (false discovery rates (FDRs), interaction term). The analysis was carried out in the window immediately upstream of m^6^A/A for all U-rich motifs previously found to be enriched around m^6^A in ECT2 targets (Arribas-Hernández et al., 2021b) and motifs previously found to be associated with m^6^A-seq peaks and ECT2-FA-CLIP peaks (Luo et al., 2014; Wei et al., 2018; Hu et al., 2021; Mao et al., 2022; Artz et al., 2023; Bian et al., 2024; Chen et al., 2024; He et al., 2024). Well-defined motifs (DRACH/RRACH, GGAU, UGUAWY and URUAY) are highlighted in bold. DRACH-like motifs are colored in purple and URUAY-like motifs in blue to facilitate their identification.

### The enrichment of URUAY/UGUA-type motifs around m^6^A sites in ECT2/ECT3 targets is largely explained by PUF targeting

Because of the substantial overlaps between PUM2/5/6 and ECT2/3 mRNA targets, we investigated further whether the enrichment of the URUAY/URUA-type motifs around m^6^A sites in ECT2/ECT3 target mRNAs may be related to their targeting by PUF proteins. We first re-analyzed the recently published comprehensive single-nucleotide-resolution m^6^A data obtained by m^6^A selective allyl chemical labeling and sequencing (m^6^A-SAC-seq) (Wang et al., 2024) to query whether there is any evidence for preferential methylation of URUAY/URUA-type motifs in Arabidopsis seedlings. This analysis demonstrated that in contrast to the previously described DRACH and GGAU methylation motifs, methylation of URUAY/URUA motifs is no higher than in random adenosines within the same set of 3’-UTRs (Fig. 3C, Supplementary Fig. S6C). This result is consistent with previously published analyses based on Nanopore direct RNA sequencing of Arabidopsis RNA (Parker et al., 2020; Arribas-Hernández et al., 2021b). Thus, URUAY/URUA is not a methylation site in Arabidopsis.

We then asked whether PUM targeting might drive the enrichment of URUAY/URUA-type motifs in the vicinity of m^6^A in ECT2/ECT3 target mRNAs. To this end, we analyzed the occurrence of URUAY around m^6^A sites in two subsets of ECT2/ECT3 target mRNAs: those targeted by either PUM2, PUM5 or PUM6 (PUM^+^), and those for which there is no evidence for PUM binding from the HT analyses (PUM^-^). We then defined 50-nt windows around m^6^A sites determined by m^6^A-SAC-seq (Wang et al., 2024) and assessed the percentage of sequences that contained one or more URUAY motifs (Fig. 3D). As a control, we performed the same analyses centered on random As instead of m^6^A sites in the 3’-UTRs of the same sets of transcripts. These analyses showed that the PUM^-^ set was significantly more strongly depleted in URUAY compared to the PUM^+^ set around m^6^A sites than around random As (Fig. 3D). Similar results were found when we considered together several PRE-like motifs (URUAY, UGUAWY, UGUAH, UGUA, URUGUAY) previously reported to be enriched around m^6^A sites in different plant species (Luo et al., 2014; Wei et al., 2018; Hu et al., 2021; Mao et al., 2022; Artz et al., 2023; Bian et al., 2024; Chen et al., 2024; He et al., 2024) (Supplementary Fig. S6D). Importantly, no other U-rich motif previously found to be enriched around m^6^A in ECT2/ECT3 targets (Arribas-Hernández et al., 2021b) showed a similar depletion in the PUM^-^ subset specifically around m^6^A sites, as measured by the significance of the difference in the motif depletion in the PUM^-^ subset comparing m^6^A-centered and random A-centered windows (Fig. 3E). We conclude that URUAY is not an m^6^A methylation motif in Arabidopsis, and that the extensive overlap in m^6^A-ECT2/3 and PUM2/5/6 mRNA targets provides an explanation for most of the observed URUAY enrichment around m^6^A sites in ECT2/3 targets. Our analyses do not, however, establish that the target overlap between PUF and ECT proteins is the exclusive reason for this enrichment as other classes of URUAY/UGUA-binding RBPs co-occurring with m^6^A-ECT may await discovery.

### A hypothesis for opposing action of PUFs and m^6^A-ECTs in controlling cellular proliferation in organ primordia

The observations of overlapping expression patterns of PUM2/6 with ECT2/3 in mitotically active cells of organ primordia (Fig. 1), as well as significantly overlapping mRNA target sets (Fig. 3B), indicate that the biological functions of PUFs and m^6^A-ECT2/3 may indeed be linked. Furthermore, since PUM5 exhibits highest expression in the RAM elongation zone characterized by cell cycle exit, yet maintains a significant overlap in mRNA targets with ECT2/3 known to stimulate cell proliferation (Arribas-Hernandez et al., 2020), the results suggest that PUFs and m^6^A-ECT may influence cellular proliferation in opposite ways. These considerations prompted us to address three additional questions. (1) Do PUFs repress growth and organogenesis? (2) If so, is there evidence for functional links to growth-stimulation by the m^6^A-ECT axis? (3) If so, do such functional links involve competitive target mRNA binding *in vivo*? The remainder of this report focuses on providing answers to these three questions.

### Increased PUM2 dosage represses growth

Since neither single T-DNA insertion mutant alleles in the *PUM2*, *PUM3, PUM5* and *PUM6* genes, nor selected double mutants (*pum2 pum6*, *pum3 pum6, pum5 pum6*) exhibited easily discernible phenotypes (Supplementary Fig. S7), we decided to probe *PUM* gene function in growth and organogenesis by increasing the dosage in mitotically active cells, taking care to avoid ectopic expression. Although results obtained by increased dosage must be interpreted with care, the precedents for dosage sensitivity of mammalian Pum2 (Gennarino et al., 2015; Gennarino et al., 2018) support the validity of the strategy. Hence, we used the strong primordium-specific promoter of the ribosomal protein-encoding gene *uS7y* (AT3G11940, formerly *RPS5A* (Weijers et al., 2001)) to drive increased expression of *PUM2-Venus* in the same expression domain as endogenous *PUM2* (*PUM2-Venus^OX^*), and built comparable *Pro_uS7y_:Venus* lines (*Venus^OX^*), all in the Col-0 ecotype, to serve as control (Fig. 4A, top row).

**Figure 4.**
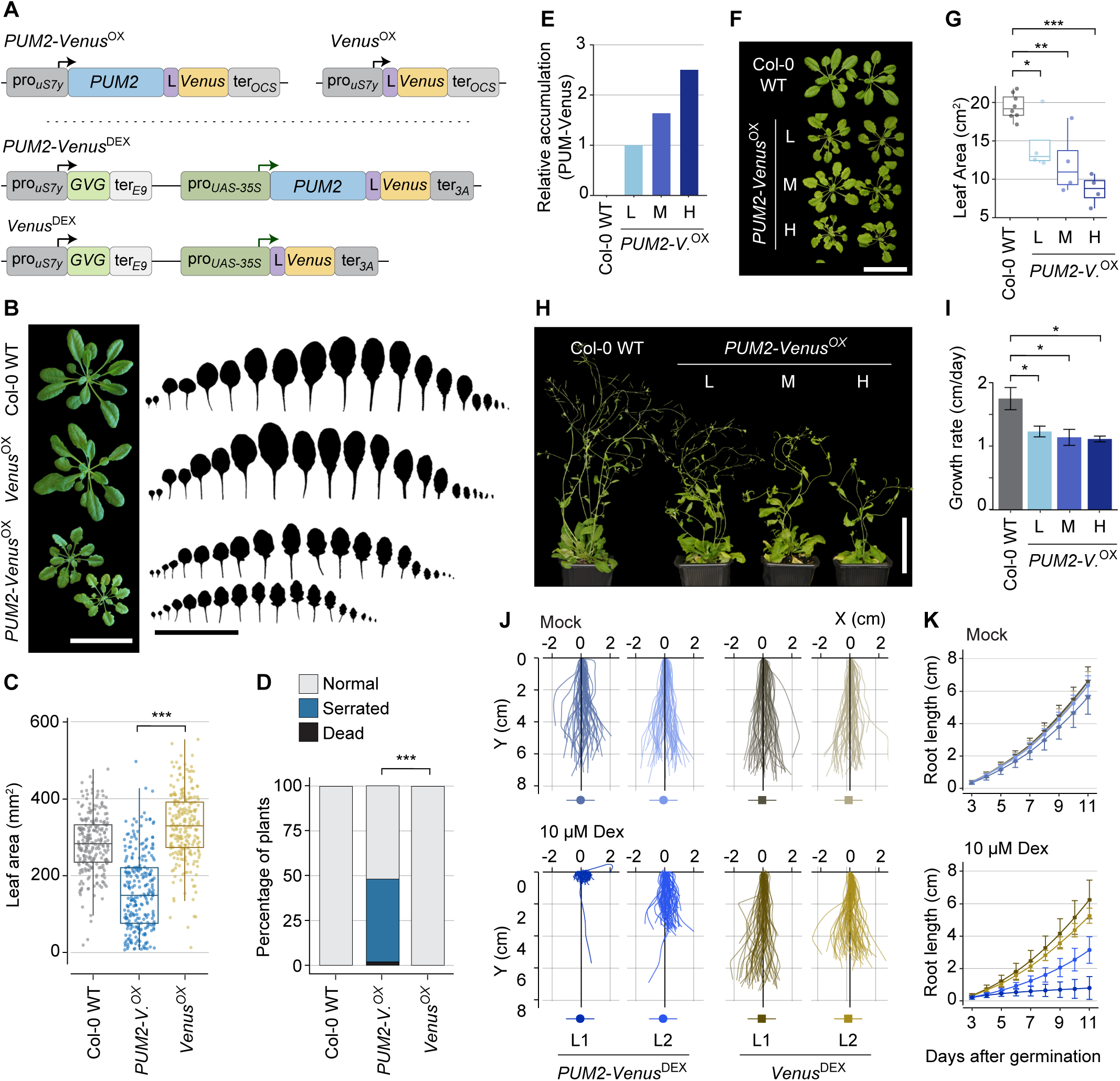
PUM2 overexpression in organ primordia represses leaf, shoot and root growth. **(A)** Schematic overview of constructs directing strong, primordium-specific constitutive (top) and dexamethasone-inducible (bottom) *PUM2-Venus* and *Venus* expression. GVG, GAL4-VP16-GR. **(B)** Leaf profiles of 4-week-old T1 plants overexpressing *PUM2-Venus* or *Venus* in organ primordia compared to those of Col-0 wild type. Scale bars, 5 cm. **(C)** Quantification of the rosette leaf area of 2.5-week-old T1 plants of the genotypes described in (B). Around 120 individual plants were examined for each genotype. ***, p-adj < 0.001 (t-test with Benjamini-Hochberg correction). **(D)** Percentage of T1 plants with the indicated leaf morphology. ‘Dead’ indicates that the plant died before phenotyping. ***, p-adj < 0.001 for Chi-square test with Benjamini-Hochberg corrections. **(E)** Quantification of protein blot analysis (Supplementary Fig. S8B) assessing the relative accumulation of PUM2-Venus in 8-day-old T2 seedlings of independent transgenic lines. Lines L (low), M (medium) and H (high) are named after their relative PUM2-Venus abundance. Line L is used as reference for the quantification. **(F-G)** Rosettes (F) and quantification of the total leaf area (G) of 4-week-old T2 plants of the lines described in (E). Scale bar, 5 cm. *, p-adj < 0.05; **, p-adj < 0.01; ***, p-adj < 0.001 for t-test with Benjamini-Hochberg correction. **(H)** 7-week-old T2 plants of the lines described in (E), showing shoot growth. Scale bar, 5 cm. **(I)** Average growth rate of the main stem in the T2 lines described in (E) during days 7^th^ to 17^th^ after bolting. Error bars indicate standard error of the mean stem growth rate. *, p-adj < 0.05 for t-test with Benjamini-Hochberg correction. **(J-K)** Representation (J) and quantification (K) of the root growth of the indicated transgenic lines from 0 to 11 days after germination, with or without dexamethasone (Dex)-induced overexpression of *PUM2-Venus* or *Venus*. Error bars in (K) indicate standard error of the mean root length.

We used two strategies to assess the effect of increased PUM2 dosage in primordial cells on leaf growth and morphology. First, we examined phenotypes of large numbers of primary transformants (T1s). Compared to the *Venus^OX^*and Col-0 wild type controls, *PUM2-Venus^OX^* lines exhibited reduced rates of leaf formation, the leaves were smaller with serrated margins (Fig. 4B to D), and stem growth was compromised (Supplementary Fig. S8C,D). Second, we selected three independent lines expressing low (L), moderate (M), and high (H) levels of PUM2-Venus according to fluorescence intensity and signal intensity in protein blots of total lysates from whole seedlings (Fig. 4E, Supplementary Fig. S8A,B), and analyzed the same traits in the second transgenic generation (T2). Similar to the observations in T1, T2 plants of the selected *PUM2-Venus^OX^* lines had smaller leaves with serrated margins, a phenotype that was more pronounced in lines with higher PUM2-Venus expression (Fig. 4E-G). The same trend was observed regarding the growth of the main stem: delayed initiation, decreased growth rate, and shorter final height (Fig. 4H,I, Supplementary Fig. S8E,F).

Since ∼2% of the primary transformants exhibited strongly stunted growth and did not grow beyond the rosette stage (Fig. 4D), their phenotypes could not be characterized in more detail. To overcome this limitation, we constructed lines expressing dexamethasone-inducible *PUM2-Venus* in the *uS7y* expression domain (Fig. 4A, bottom). When such lines were grown in continuous presence of the inducer, severe, in some cases nearly total, growth restriction as quantified by root growth rates was observed, and the degree of root growth inhibition correlated with the PUM2-Venus expression level (Fig. 4J,K, Supplementary Fig. S8G). We conclude that increased PUM2 dosage in organ primordia leads to growth restriction and altered leaf morphology, reminiscent of the effect of simultaneous inactivation of *ECT2*, *ECT3* and *ECT4* (Arribas-Hernández et al., 2018).

### A PUM2 mutant mimicking constitutive TOR-dependent phosphorylation is a less potent growth repressor

Although dosage manipulation is a valid test of gene function, it is prone to misinterpretation because artificial effects acquired under unnaturally high concentrations may contribute to the observed phenotypes. While it is difficult to entirely exclude this possibility, controls with point mutants expected to modulate the functionality of the protein are helpful to corroborate the validity of the conclusions drawn on the basis of overexpression phenotypes. We reasoned that the identification of PUF phosphorylation sites dependent on the major growth activator TOR kinase (Van Leene et al., 2019) would be of use in this context. Specifically, since TOR activity stimulates the growth program (Xiong and Sheen, 2014; McCready et al., 2020), the expectation would be that TOR-dependent phosphorylation of PUFs should alleviate their potential as growth repressors. To test this prediction, we constructed PUM2 point mutants with all four Ser/Thr residues subject to TOR-dependent phosphorylation changed to either Glu (mimicking constitutive phosphorylation, “phospho-mimic”, “PUM2^4A^”) or Ala (mimicking constitutive dephosphorylation, “phospho-dead”, “PUM2^4E^”) (Fig. 5A), and conducted leaf area measurements in primary transformants grown side-by-side. These constructs were introduced in a CRISPR-Cas9-induced, T-DNA-free *ect2* knockout mutant background (*ect2-6*) that does not have conspicuous growth defects (Supplementary Fig. S9), as described previously for other *ect2* knockout alleles (Arribas-Hernández et al., 2018). The results showed that the growth repression conferred by the PUM2 phospho-mimic mutant was significantly reduced compared to the wild type and phospho-dead mutant (Fig. 5B-E). These observations corroborate the conclusion that growth repression is a wild type function of PUM2, and suggest that it acts in the context of TOR-controlled growth.

**Figure 5.**
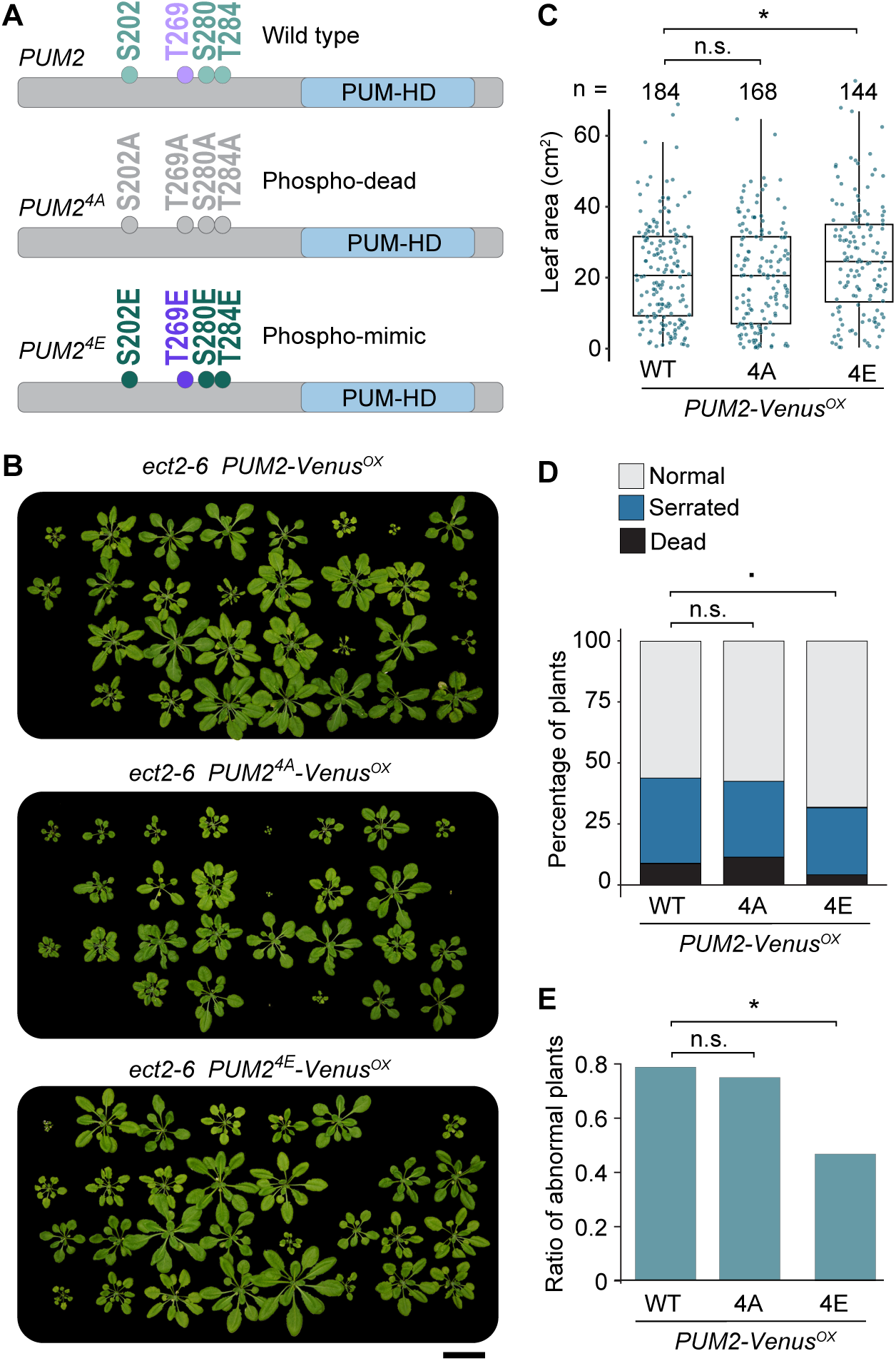
TOR-mediated phosphorylation of the PUM2 IDR alleviates PUM2-mediated growth repression. **(A)** Schematic representation of PUM2 wild type (PUM2^WT^) and PUM2 phosphosite mutants (PUM2^4A^ and PUM2^4E^). The Ser/Thr phosphorylation sites at the N-terminal IDR of PUM2 are dependent on either TOR (green) or both TOR and MAP kinases (purple). The PUM homology domain (PUM-HD) is highlighted in blue. **(B)** Examples of 4.5-week-old *ect2-6* plants (Supplementary Fig. S9) overexpressing either PUM2 wild type or PUM2 phosphosite mutants. Scale bar, 5 cm. **(C)** Quantification of the rosette size of 3-week-old T1 plants expressing PUM2 wild type or PUM2 phosphosite mutants. Statistical significance was assessed using Student’s t-test. **(D-E)** Analyses of T1 plants overexpressing PUM2 wild type or PUM2 phosphosite mutants. (D) Percentages of plants with the indicated leaf morphology phenotype. ‘Serrated’ indicates serrated leaf edges, and ‘dead’ indicates that the plant was dead at the time of phenotyping. (E) Percentages of abnormal plants (abnormal/normal). In D,E, statistical significance was assessed using Chi-square test. For C-E: n.s., not significant;., p < 0.1; *, p < 0.05; **, p < 0.01.

### PUM2 overexpression confers defective pollen development

In addition to the growth phenotypes characterized above, we noticed that *PUM2-Venus* overexpressing plants showed greatly reduced fertility (Fig. 6A-B), probably because of defective pollen formation, since almost no shedding of pollen grains was observed by visual inspection (Fig. 6C). Indeed, Alexander staining for mature pollen verified that PUM2-Venus-overexpressing plants produced very few mature pollen grains (Fig. 6D). We note that several of the PUM1-PUM6 proteins are expressed in the microspore, bicellular and tricellular stages of pollen formation (Supplementary Fig. S1C), indicating that the effect of Pro*_uS7y_*-driven *PUM2-Venus* expression relies on increased dosage of a PUM-function already present in these cells, not on ectopic expression.

**Figure 6.**
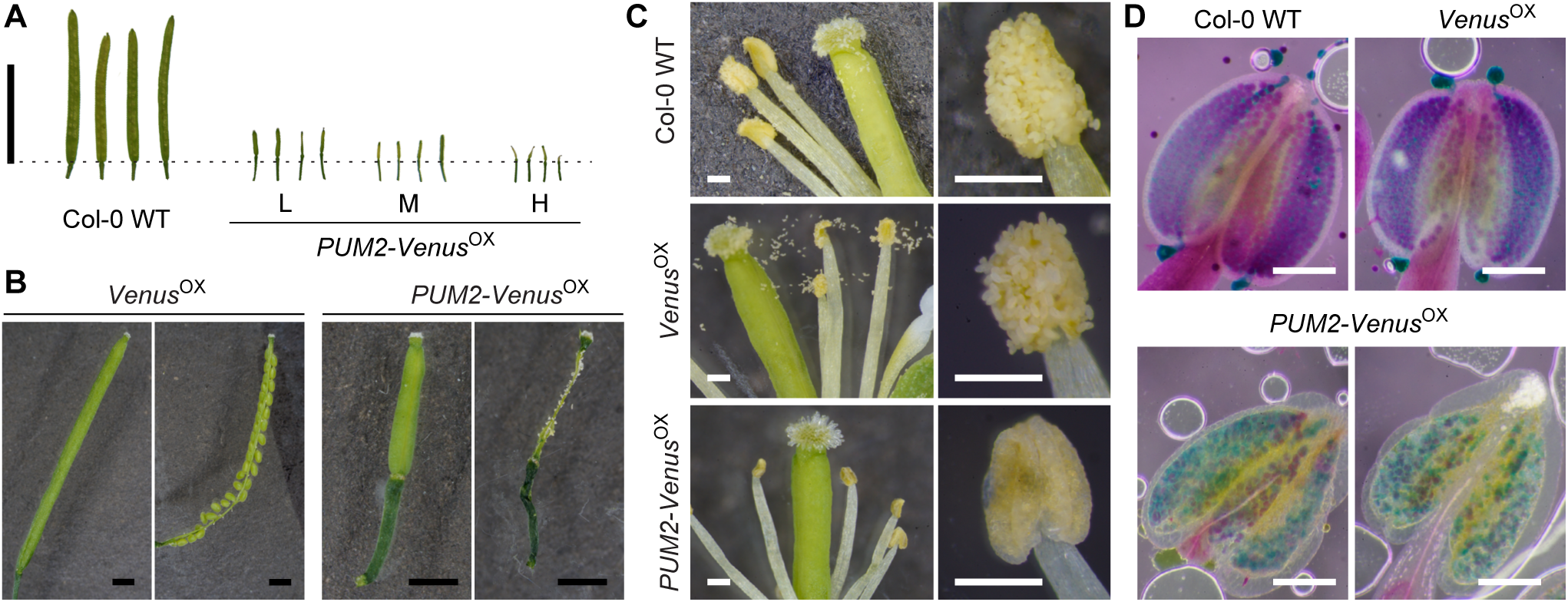
PUM2 overexpression causes defective pollen development. **(A-B)** Silique characterization of T2 plants overexpressing *PUM2-Venus* (*PUM2-Venus^OX^*) compared to Col-0 wild type (WT) or transgenic control (*Venus^OX^*). (A) Appearance of several siliques. Scale bar, 1 cm. (B) Silique dissection. Scale bars, 1 mm. **(C)** Pistil and stamens of stage 14-15 flowers of the genotypes in (A). Scale bars, 0.2 mm. **(D)** Anthers of stage 12 flowers from the genotypes described in (A) subjected to Alexander staining. Viable pollen is stained in red-purple colors, and non-viable pollen turns green-blue. For *PUM2-Venus^OX^*, two independent transgenic lines are shown. Scale bars, 0.1 mm.

### Synthetic effects of PUM2 overexpression and ECT2 or ECT2/ECT3/ECT4 inactivation

We next compared the effect of PUM2 overexpression in wild type and *ect* mutant backgrounds to judge synthetic effects that may reveal mechanistic links between PUM-and m^6^A-ECT-mediated gene regulation. To this end, we introduced the *Pro_uS7y_:PUM2-Venus* transgene into the T-DNA-free *ect2-6* mutant to avoid possible T-DNA interference (Mlotshwa et al., 2010) with previously characterized *ect2* T-DNA alleles (Arribas-Hernández et al., 2018; Scutenaire et al., 2018; Wei et al., 2018; Arribas-Hernandez et al., 2020). Two independent *ect2-6 PUM2-Venus^OX^* lines with moderate or strong growth phenotype (Fig. 7A,B) were then crossed to either wild type Col-0 or to *ect2-6*, to generate F1 plants with PUM2-Venus overexpression from the same *Pro_uS7y_:PUM2-Venus* transgene either in the presence of ECT2 protein (*ect2-6/+* heterozygotes resulting from the cross to Col-0), or in the absence of ECT2 protein (*ect2-6* homozygotes resulting from the cross to *ect2-6*). For both lines, we observed markedly stronger growth repression as measured by leaf area when *PUM2-Venus* was overexpressed in the absence of ECT2 than in its presence (Fig. 7C). This genetic interaction indicates mechanistic links between growth control mediated by PUM2 and ECT2, consistent with their co-expression and overlapping mRNA target sets.

**Figure 7.**
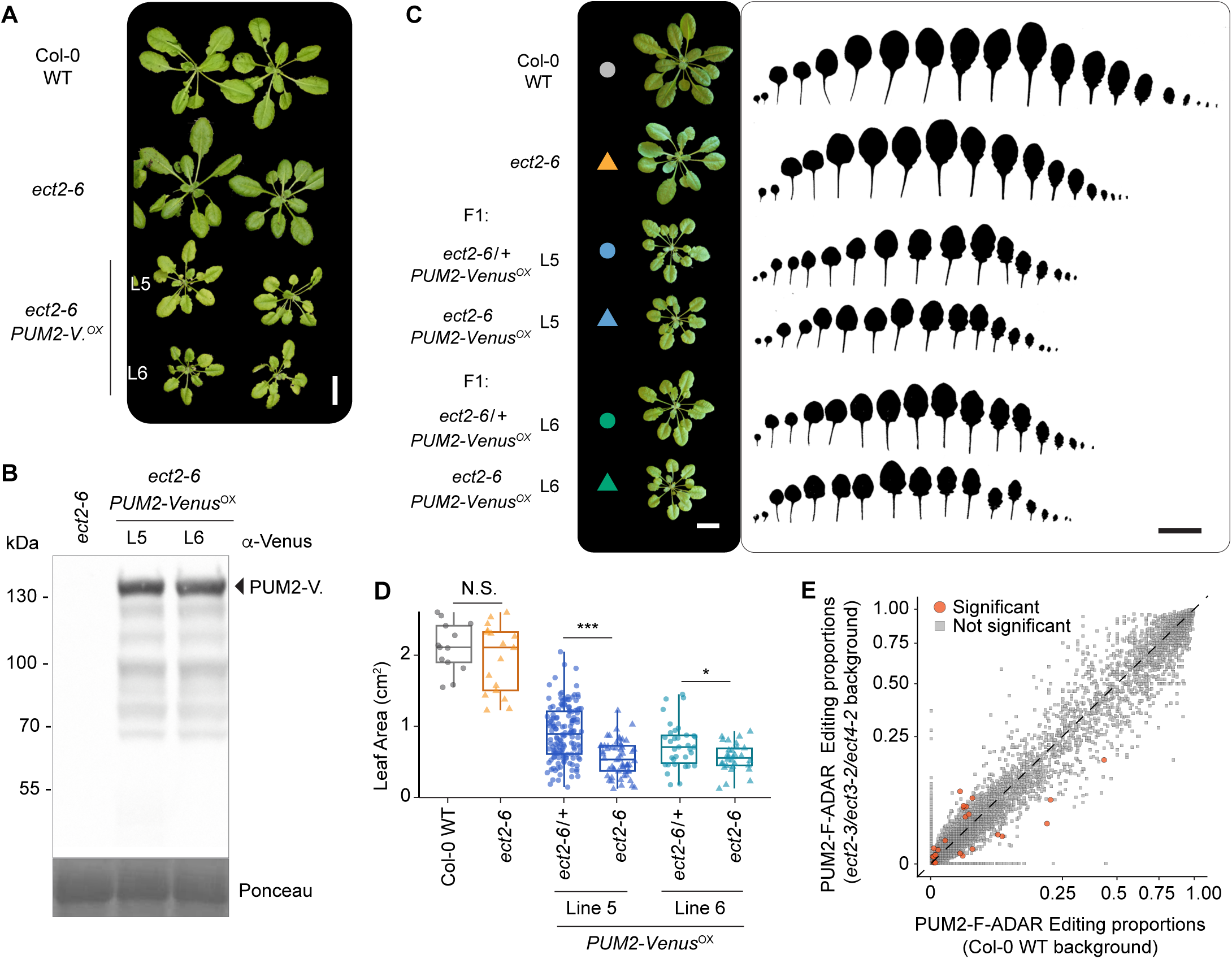
Evidence for a genetic interaction between *PUM2* and *ECT2*. **(A)** Representative 4-week-old T2 plants of two independent *ect2-6 PUM2-Venus*^OX^ lines, and controls. Scale bar, 2 cm. **(B)** Western blot of the plants in (A), showing PUM2-Venus accumulation. **(C)** Leaf profiling of 4-week-old plants of the indicated genotypes. Scale bars, 2 cm. **(D)** Quantification of rosette size of the genotypes in (C). *, p-adj < 0.05; ***, p-adj < 0.001 (t-test with Benjamini-Hochberg correction). **(E)** Scatter plot of the editing proportions (E.P.) of PUM2-FLAG-ADAR-edited sites in the triple *ect2-3/ect3-2/ect4-2* mutant (Arribas-Hernández et al., 2020) compared to an otherwise wild type Col-0 background. Orange dots indicate sites with significantly different editing proportions between the two backgrounds.

We also wished to compare the effect of PUM2 overexpression in wild type and *ect2-3 ect3-2 ect4-2* ((Arribas-Hernandez et al., 2020), henceforth *ect2/3/4*) mutants. Due to the presence of three T-DNAs in the *ect2/3/4* mutant and to avoid silencing of the *PUM2-Venus* transgene over generations, we chose to conduct this test by direct analysis of leaf formation, size and morphology reproduction defects in large numbers of independent transgenic lines in the first transgenic generation. We observed a clear exacerbation of leaf morphology defects, delay of leaf formation and reduction of leaf size in PUM2-Venus-expressing *ect2/3/4* mutants (Supplementary Fig. S10 A to D). The proportion of plants with defective pollen development was also clearly higher when PUM2-Venus was overexpressed in the *ect2/3/4* background compared to the wild type Col-0 background, despite the fact the *ect2/3/4* on its own showed no detectable pollen defects (Supplementary Fig. S10 E). These results corroborate the conclusion that a defect in the m^6^A-YTHDF system enhances the effect of PUM2 overexpression, indicating mechanistic links between mRNA control exerted by the PUF and YTHDF families.

### ECTs do not appear to compete with PUMs for mRNA target binding in vivo

A simple molecular model to account for the synergistic interaction between *PUM2* overexpression and *ECT2* inactivation would be competitive binding to their mRNA targets. In that case, PUM2 occupancy would increase in the absence of major ECT proteins, thereby contributing to growth repression in the *ect2/ect3/ect4* mutant background.

We previously used quantitative differences in HT editing proportions between the same ADAR fusion expressed in different genetic backgrounds to show that ALBA proteins assist ECT2 target binding *in vivo* and that ECT2 and ECT3 compete for the same targets *in vivo* (Arribas-Hernández et al., 2021a; Reichel et al., 2024). We applied the same experimental and computational strategy to test whether the presence of ECT2/3/4 influences PUM2 target mRNA association *in vivo*. Thus, we selected independent PUM2-FLAG-ADAR lines with comparable transgene expression in wild type and *ect2/ect3/ect4* backgrounds (five lines of each type, Supplementary Fig. S11), and used them to compare editing proportions of PUM2-FLAG-ADAR-edited sites in target mRNAs in the presence or absence of ECT2/3/4. Analysis of this HT dataset showed that very few sites had differential PUM2-dependent editing between wild type and mutant plants, and no clear trend towards either background was discernible (Fig. 7F). Hence, HyperTRIBE provides no evidence for competing action of ECTs on PUM2-binding to target mRNAs *in vivo*.

### Proteins physically associated with PUM2 and ECT2 overlap only marginally

We reasoned that since PUM2 binding to mRNA targets shared with ECT2 and ECT3 was unaffected by knockout of *ECT2/3/4*, PUFs and ECTs may be associated with the same mRNA molecules, raising the possibility of physical interactions with shared sets of proteins, including association of PUFs and ECTs. To test this possibility, we immuno-affinity purified PUM2-Venus and PUM5-Venus expressed under their endogenous promoters in Arabidopsis stable lines, and identified associated proteins by liquid chromatography-mass spectrometry of tryptic peptides generated by digestion of entire immunopurified fractions (Supplementary Fig. S12). RNaseA was included in the lysis buffer to prevent association of RBPs mediated merely by RNA. These analyses uncovered two fully overlapping sets of 128 and 161 proteins enriched in the PUM2-Venus and PUM5-Venus purifications compared to the Venus control expressed under the *uS7y* promoter (Fig. 8A). RNA-binding proteins, molecular chaperones and photosynthesis-related proteins were predominant in the PUM2-associated set (Fig. 8B,C). When compared to the group of ECT2 interactors that require its N-terminal IDR for association (Tankmar et al., 2023), we only found a marginal overlap of 12 proteins, including 7 chaperones and 5 photosynthesis-related proteins (Fig. 8D and Supplementary Fig. S13A). Thus, the PUM2-associated set of proteins identified here does not immediately reveal the nature of the functional links between PUM2 and ECT2 implied by their genetic interaction.

**Figure 8.**
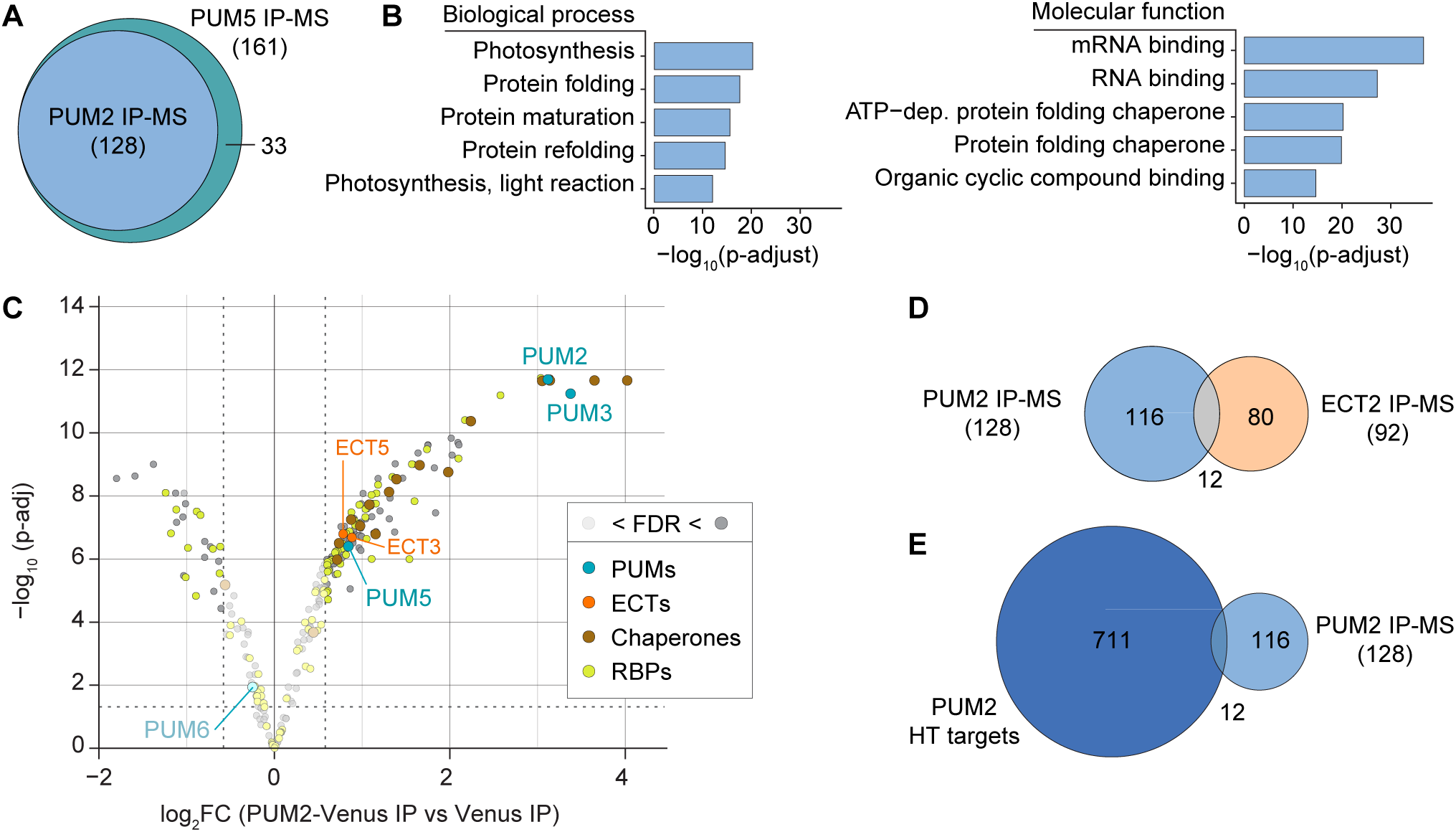
Overview of PUM2-and PUM5-associated proteins. **(A)** Venn diagram showing the overlap of PUM2- and PUM5-associated proteins identified by immunoprecipitation-mass spectrometry (IP-MS). **(B)** Gene ontology-term analysis of proteins associated with PUM2 IP-MS. The complete set of Arabidopsis proteins was used as background in the GO-term analysis. **(C)** Volcano plots showing differentially enriched proteins in PUM2-Venus IP-MS (compared to Venus), with other PUF proteins, chaperones, ECTs, and other RNA-binding proteins (RBPs) highlighted. **(D)** Venn diagram showing the overlap of PUM2- and ECT2-associated proteins (Tankmar et al., 2023) identified by IP-MS. **(E)** Venn diagram showing the overlap of PUM2 HyperTRIBE (HT) targets and PUM2 interacting proteins identified by IP-MS.

Although ECT2 was not among the RBPs significantly enriched in the PUM2-Venus immuno-purified fractions, ECT3 and ECT5 were included in this set (Fig. 8C). We hesitate to interpret this observation as evidence for physical association between PUM2 and YTHDFs bound to common target mRNAs for two reasons. First, since ECT2 is the most highly expressed YTHDF protein in the PUM2 expression domain, and its targets overlap as much with PUM2 targets as those of ECT3 (Supplementary Fig. S13B), one would expect to also find ECT2 associated with PUM2 if indeed this interaction takes place on target mRNAs. We did not detect any ECT2 peptides in the PUM2-Venus immuno-affinity purified fractions. Second, ECT3 and ECT5, but not ECT2, belong to a small set of genes whose encoded mRNAs are PUM2 HT targets and whose encoded proteins are enriched in PUM2 immunopurified fractions (Fig. 8E, Supplementary Table 5 and Supplementary Fig. S13C). Hence, the exclusive identification of ECT3 and ECT5 might result from PUM2 interactions with the nascent ECT3 and ECT5 proteins, mediated by PUM2 binding to the ECT3 and ECT5 mRNAs. We conclude that the PUM2 IP-MS data provides no clear evidence for a physical PUM2-YTHDF interaction at shared target mRNAs, although the data is clearly not sufficient to exclude such an interaction.

## DISCUSSION

### Plant PUF function in the context of m^6^A-YTHDF-mediated growth control

The PUF proteins are among the most intensely studied mRNA-binding proteins owing to their deep conservation and important biological functions in many eukaryotic organisms. Despite the discovery of key functions of Pumilio in Drosophila and FBF in *C. elegans* more than 30 years ago, only limited progress has been made to uncover the molecular and biological functions of PUF proteins in plants. This study offers advances to this end in three closely connected ways. First, it identifies mRNA targets *in vivo* of PUM2, PUM5 and PUM6 transcriptome-wide, and shows that the target overlap with ECT2/3 is substantial. Second, it provides evidence that PUM2 promotes growth repression and that PUM2/6 and ECT2/3 are co-expressed in mitotically active cells of root and shoot meristems. Third, it demonstrates genetic interaction between *PUM2* and *ECT2/ECT3*. Taken together, these results allow the formulation of a working model for biological function of canonical plant PUF proteins in the context of growth control: PUF proteins and m^6^A-YTHDF target a common set of mRNAs in organ primordia and exert opposite effects on growth, with PUFs acting to repress and m^6^A-YTHDF acting to stimulate it.

We stress that while this model is clearly consistent with the observations reported here, not all aspects of it can be regarded as fully proven. This is because the tests of biological functions of PUF proteins reported here rely on analyses of consequences of increased dosage of PUM2 in its natural expression domain. While the literature is rich in examples where phenotypes conferred by increased dosage successfully reveal the wild type gene function, it is difficult to judge whether observed phenotypes really are exclusively due to enhanced natural function of the overexpressed protein, or whether the increased protein concentration allows artificial neo-function that may cause or contribute to the phenotypes. For example, in the case of PUM2, increased concentration may allow binding of PUM2 to mRNAs not targeted naturally by PUM proteins. We note, however, that the effect on growth and organogenesis is consistent not only with the expression pattern of PUM2, PUM5 and PUM6, but also with the significant mRNA target overlap with ECT2 and ECT3 that clearly promote growth by stimulating proliferation of primordial cells (Arribas-Hernández et al., 2018; Arribas-Hernandez et al., 2020). In addition, the use of the PUM2 point mutant mimicking constitutive phosphorylation dependent on the key growth activator TOR provides corroborating, if still circumstantial, evidence that growth repression is a function of the PUM2 wild type protein. The PUM2 phosphomimic mutant was clearly less potent as a growth repressor than wild type, consistent with the well-established growth-promoting function of TOR.

We note that our study does not reveal the mechanism by which PUFs repress growth, including the mechanistic underpinnings of the functional interplay with m^6^A-YTHDF. The existence of such interplay is best supported by the synthetic effects of *PUM2* overexpression and *ECT2* inactivation. The observed synergy indicates mechanistic coupling, particularly because inactivation of *ECT2* alone only has very subtle phenotypic consequences, limiting the possibility that the synthetic interaction with PUM2 overexpression is caused by indirect effects of the inactivation of *ECT2*. How this coupling works is now a question of great interest. We disfavor the simplest model of competitive target binding, because the editing proportions conferred by PUM2-FLAG-ADAR and PUM6-FLAG-ADAR on target mRNAs *in vivo* were unaltered upon knockout of *ECT2*, *ECT3* and *ECT4*. Thus, overexpression may simply increase the fraction of mRNA targets co-bound by both PUM2 and YTHDFs, tipping the balance in favor of PUM2-mediated growth repression. It is also possible that PUM2 overexpression saturates enzymes involved in its post-translational modification such that overexpression changes the mRNA-bound PUM2 population from predominantly modified to predominant unmodified, perhaps exacerbating repressive activity. This possibility would explain why overexpression of both wild type and phospho-dead PUM2 conferred equally strong growth repression, while the phospho-mimic mutant was significantly less potent. In any case, our results indicate that the binding of PUFs and YTHDFs to their common target mRNAs occurs independently, despite their linked biological function. This scenario is reminiscent of the functions of Drosophila Pumilio and the RNA-binding protein Brain Tumour (Brat) in the regulation of the *hb* mRNA (Loedige et al., 2014; Arvola et al., 2017).

Given that the direct interaction between ECTs and cytoplasmic poly(A) binding proteins (PABPs) (Song et al., 2023; Tankmar et al., 2023) is of key importance for organogenesis in Arabidopsis (Tankmar et al., 2023), and that regulatory functions of yeast and metazoan PUF proteins involve interactions with PABP (Nakahata et al., 2001; Padmanabhan and Richter, 2006; Chritton and Wickens, 2011; Campbell et al., 2012; Weidmann et al., 2014; van de Poll et al., 2023), one may anticipate that the functions of PABPs could be key to the PUM-YTHDF interplay. Nonetheless, our immuno-precipitation-LC-MS analyses of proteins co-purifying with PUM2-Venus and PUM5-Venus in the presence of RNase did not find PABPs to be significantly enriched over the Venus background, suggesting that any interaction in the context of an intact mRNP may be too weak to be detected upon dilution and mRNA cleavage.

### A function of PUMs in pollen development

In addition to growth inhibition, we found that the most striking effects of increased PUM2 dosage was strongly reduced efficiency of pollen maturation. We did not clearly define the stage at which pollen maturation fails in PUM2 overexpressing plants, but note that an involvement of PUFs in development of germ cells is in line with the biological functions described in animals, as is the control of stem cell proliferation (Wickens et al., 2002). Hence, more broadly, multicellular life forms use PUF proteins to exert control over similar fundamental aspects of their life cycles in both animals and plants, and it will be of interest to understand whether this control involves recurrent or divergent molecular mechanisms.

### A molecular explanation for the enrichment of URUAY around m^6^A sites in Arabidopsis

A particularly important insight deriving from the identification of the PUF mRNA target sets concerns the puzzling enrichment of the URUAY motif around m^6^A sites in different plant species, interpreted by some as evidence that URUAY is a plant-specific methylation motif (Wei et al., 2018). Our reanalysis of the comprehensive single-nucleotide-resolution m^6^A maps obtained through m^6^A-SAC-seq (Wang et al., 2024) clearly demonstrates that URUAY is not, as a matter of fact, a methylation motif in Arabidopsis. Indeed, the URUAY enrichment around m^6^A in ECT2/ECT3 targets largely disappears upon removal of PUF mRNA targets from ECT2/ECT3 target sets, effectively demonstrating that functional links between PUFs and m^6^A-YTHDF manifested by their shared target sets underlies most of the URUAY enrichment around m^6^A sites in plants.

## MATERIALS AND METHODS

### Plant material

All lines used in this study are of the *Arabidopsis thaliana* Colombia-0 (Col-0) ecotype and are listed in Supplementary Table S1. T-DNA insertion lines were obtained from Nottingham Arabidopsis Stock Centre. Transgenic lines and CRISPR-Cas9-engineered mutants generated in this study are further described in subsequent sections.

### Growth conditions

For seed germination on Murashige & Skoog (MS)-agar plates, Arabidopsis seeds were first surface-sterilized in a laminar flowhood, where seeds were treated with 3 mL 70% ethanol for 2 min, followed by 6 mL 1.5% NaOCl/0.05% Tween-20 for 10 min, and two times washing with autoclaved Milli-Q water. A small amount of melted 0.8x MS-agar media (4.4 g/L MS-Basal Salt (PhytoTech Labs), 1 g/L sucrose, and 0.8 g/L Bacto agar) was added to the falcon tube to suspend the seeds, and seeds were plated on 1xMS–agar plates supplemented with antibiotics for transgene selection or chemicals for induction of gene expression. The plates were sealed with 3M microporous surgical tape, and cold-stratified at 4°C in darkness for 2-5 days. After cold stratification, the plates were transferred to Aralab growth chamber (Fitoclima 600) with a temperature of 21-22°C, light intensity of ∼120 µmol m^-2^ s^-1^, and 16 hr light/ 8 hr darkness photoperiod. After 6-12 days growth on MS-agar plates, seedlings were transferred to soil (Plugg/Såjord [seed compost], SW HORTO A/S) containing 4% perlite, 4% vermiculite and 1% slow-release fertilizer (Multicote™ Agri), and grown in greenhouses for seed production or Aralab growth chamber with a 16 hr light / 8 hr darkness photoperiod, light intensity of ∼120 µmol m^-2^ s^-1^, and a temperature of 21-22°C day/16-17 °C night for phenotypic characterization.

### CRISPR-Cas9 gene editing in Arabidopsis

CRISPR-Cas9 gene editing was performed as described (Tsutsui and Higashiyama, 2017). Guide RNAs were designed with CCTop (Stemmer et al., 2015). Two complementary oligonucleotides were annealed and ligated into pKIR using T4 DNA ligase. Constructs encoding two guide RNAs with target sites 300-500 bp apart were then transformed into *Arabidopsis thaliana* plants using floral dip (Clough and Bent, 1998). T1 plants were antibiotic-selected, genotyped for the presence of deletions and sequenced. Cas9-free T2 plants were selected by genotyping and examining red fluorescence of seeds, and plants were sequenced again to make sure that the mutation was inherited from T1 plants. T3 plants were selected for carrying homozygotic mutations. The oligonucleotides used to build CRISPR constructs can be found in Supplementary Table S2.

### Genotyping of T-DNA insertion lines and *ect2* CRISPR mutant (*ect2-6*)

Genotyping was performed by two distinct PCR-based methods. In the first, genomic DNA from young leaves was isolated using a urea-sarcosyl lysis buffer, followed by phenol-chloroform purification, and used as templates for genotyping PCR. In the second, lysates of young leaves without further purification were directly added to the Thermo Scientific Phire Hot Start II PCR reaction. T-DNA lines with the corresponding PCR primers for genotyping are listed Supplementary Table S2.

### Generation of PUM-Venus and Venus transgenic lines

Recombinant DNA constructs were generated by USER cloning, as described (Bitinaite and Nichols, 2009; Arribas-Hernandez et al., 2020). Briefly, gene fragments were PCR-amplified using dU-containing primers (TAGC Copenhagen) and KAPA DNA polymerase (Roche). The purified USER-cloning gene fragments were thenligated into linearized vectors containing a USER-cassette. Specifically, two USER-cassette containing vectors were built for this study. The pLIFE5s vector was modified from pLIFE5. It is generated by PacI-mediated restriction cloning, replacing the phosphinothricin resistance gene with the sulfadiazine resistance gene in pLIFE5. The pTA7002UR vector was modified from pTA7002U. First, site-directed mutagenesis was performed by QuickChange (Agilent) to remove the PacI restriction site within the *us7y* promoter sequence in plasmid pGGA012. The PacI-site-free *us7y* promoter sequence was then PCR-amplified and inserted in PTA7002U to replace the 35S promoter driving GAL4-VP16-GR expression with PacI restriction cloning. Primers, PCR templates, and corresponding vectors used for USER cloning in this study are listed in Supplementary Table S2.

After ligation of USER fragments and lineralized vector, the constructs were electroporated into *Agrobacterium tumefaciens* GV3101 and the resulting strains were used to transform Arabidopsis by floral dip (Clough and Bent, 1998). Selection of primary transformants (T1) was done on MS-agar plates supplemented with the appropriate antibiotics. For each genotype of plants, multiple T2 lines with single-locus insertions were selected by inspecting the segregation on antibiotic-supplemented plates.

### Total protein extraction and western blot

Fresh plant tissue was collected in liquid nitrogen and ground with either mortar-pestle or by two minutes of vigorous shaking with metal beads in a plant tissue homogenizer at 30 hz, with 30 sec intervals where tubes were cooled in liquid nitrogen to avoid tissue thawing. Three volumes (v/w) of cold IP buffer (50 mM Tris-HCl pH 7.5, 150 mM NaCl, 10% glycerol, 5 mM MgCl_2_, and 0.1% Nonidet P40), freshly supplemented with protease inhibitor (Roche Complete tablets) and 1 mM DTT, was added to the ground tissue, immediately followed by vigorous vortexing. The lysate was centrifuged at 13000 x *g* for 10 min at 4°C, and the supernatant was transferred to clean, pre-cooled Eppendorf tubes, mixed with LDS sample buffer (277.8 mM Tris-HCl pH 6.8, 44.4% (v/v) glycerol, 4.4% LDS, and 0.02% bromophenol blue) to a final concentration of 1x LDS, and denatured at 75 °C for 10 min. Protein samples were loaded on a 4-20% Criterion TGX Precast gel (Biorad) with PageRuler protein ladder (Thermo Scientific) to indicate the size of proteins. The gel was run in 1x Running Buffer (25 mM Tris, 192 mM Glycine, 0.1% SDS) at 100V for ∼2 hr on ice. After electrophoresis, proteins were wet-transferred to Protran Premium nitrocellulose membrane (Cytiva) in cold transfer buffer (25 mM Tris, 192 mM glycine, 20% ethanol) at 80 V for 1 hr on ice, followed by Ponceau staining. The membrane was then blocked in 5% PBST-milk (1x PBST buffer (137 mM NaCl, 2.7 mM KCl, 10 mM Na_2_HPO_4_, 1.8 mM KH_2_PO_4_, 0.05% Tween-20, pH 7.4), with 5% (w/v) skimmed milk powder) for 30 min at room temperature. After blocking, membranes were incubated with primary antibody in 5% PBST-milk at 4° C overnight. The following day, membranes were washed 3 times with 1x PBST buffer for 5 min each, followed by 90 min room-temperature incubation with HRP-conjugated secondary antibody in 5% PBST-milk, except for FLAG for which the primary antibody (Sigma, F3165) was HRP-conjugated. After the incubation the membrane was further washed with 1x PBST buffer 3 times 5 min each, and MilliQ water 2 times 5 min each. Supersignal West Femto (Thermo Scientific) or enhanced chemiluminescence (ECL, Thermo Scientific) was added to the membrane and incubate in darkness for ∼1 min for signal development, and pictures of the membrane were taken in the darkroom with a SONY ILCE-7S camera controlled by Capture One 11. The western blot images were analyzed with Fiji (Schindelin et al., 2012) to quantify signal intensities when necessary.

### Total RNA extraction

Fresh plant tissue was collected and ground as described in the previous section. 1 mL of TRI Reagent (Sigma) was added to 100 mg of liquid-nitrogen frozen plant tissue powder and immediately homogenized by vortexing. The samples were kept on ice unless specified otherwise, and centrifugations were performed at 4 °C at 13,000 *g*. 200 μL of chloroform was added to the mixture and vortexed. The tubes were then centrifuged for 10 min. The upper aqueous phase was carefully transferred to a new, pre-cooled Eppendorf tube with an equal volume of cold isopropanol, followed by incubation of 30 min at room temperature. The tubes were then centrifuged for 15 min to pellet precipitated RNA. The supernatant was removed with a pipette, and the pellet was washed in cold 70% ethanol twice. The pellet was then resuspended in nuclease-free water, and polysaccharides were precipitated by addition of 1/10 volume of cold 99% ethanol and 1/30 volume of 3 M NaOAc pH 5.2, followed by 30 min incubation on ice. The tubes were then centrifuged at 4°C for 15 min to pellet precipitated polysaccharides. The supernatant was transferred to a new, pre-cooled Eppendorf tube with 2.5 volume of cold 99% ethanol and 0.1 volume of 3 M NaOAc pH 5.2, followed by incubation at -80 °C overnight and centrifugation for 30 min. The RNA pellet was washed in cold 70% ethanol for twice and residual ethanol was carefully removed with a pipette. The RNA pellet was then air-dried for 5 min and resuspended in nuclease-free water. The quality of the RNA was then examined by electrophoresis or Bioanalyzer run, and the concentration was measured with Nanodrop.

### Fluorescence microscopy

PUM-Venus expression and ECT2-mCherry expression in Arabidopsis were examined by fluorescence microscopy with a stereomicroscope (Leica MZ16 F equipped with 0.63x -12x objective and filters for GFP and RFP) or a confocal microscope (Zeiss LSM700 equipped with 10x-63x objective and two photomultiplier tube (PMT) detectors). Stereomicroscope images of seedlings on MS-agar plates were taken with a SONY α6000 camera and processed with Fiji. Confocal images of living seedlings on glass slides were obtained using Zeiss Zen and further processed with Fiji.

### Plant phenotypic analyses and statistics

For all phenotypic characterization, plants were grown in parallel, i.e. plating, germination and, when required, transfer to soil took place on the same day. For stem length measurement, the main stem was pulled straight and a ruler was used to measure the length of main stem from bottom to top. For T1 plants, the stem length was measured once at 29 days after germination (DAG), and for T2 and F1 plants, the stem length was measured every 2 days from DAG 29 to DAG 67. Analysis of stem length data was done in R. The mean and standard deviation were calculated for each transgenic line. For stem length measurements recorded every 2 days, the statistics were calculated for every 2-day interval. The stem length of T2 and F1 plants were also aligned to the first day of bolting (DAB). The growth rate of the main stem was calculated from DAB 7 to DAB 17, where the growth curve was approximately linear. The mean stem length for each line was modelled as a linear function of DAB, and the slope was used as the estimated growth rate. To assess whether the stem length and growth rate significantly differed among different genotypes, *t*-tests with Benjamini-Hochberg adjustment for multiple comparisons were performed.

For measurement of rosette size, pictures of plants between 17-32 DAG on soil were taken with a SONY IL-7S camera. The camera was positioned vertically above the rosettes to capture the top view, and a ruler was included in each image. Images were then processed in Adobe Photoshop and Fiji (Schindelin et al., 2012). First, the raw pictures from camera were pre-processed in Photoshop. The ‘magic wand’ tool was used to select the green plants, and the background was masked with black to avoid the interference of perlite, vermiculite and fertilizer (with white or yellow colors) in subsequent steps. The pre-processed, RGB-colored images were converted to 8-bit binary images in Fiji, with the green plants rendered white and the background black. Overlapping plants were manually separated by drawing black lines to ensure that the software did not mistakenly identify multiple rosettes as a single object. To measure the vertical projection area of each rosette on the soil surface, a scale was set using the ruler included in each image. The ’Analyze Particles’ function in Fiji was then used to measure area, centroid coordinates (X and Y), and Feret’s diameter for each object. The measurements were imported into Excel and sorted based on the X and Y coordinates. Genotype and plant ID were manually added and the dataset was subsequently imported into R for statistical analysis. Mean and standard deviation of rosette size were calculated for each line. To assess the statistical significance of the observed differences in rosette size and stem growth between T2 lines, *t*-tests with Benjamini-Hochberg adjustment for multiple comparisons method) were performed.

The rosette morphology of plants was assessed in 3-4 week old plants. Plants were categorized as normal, serrated leaves, curly leaves, or dead. Combinations between different categories were allowed. The total number of plants with rosette morphology different from wild type was calculated per line and imported to R. To assess whether the distribution of rosette morphology changes differed significantly among plant lines, chi-squared test was performed.

For characterization of root growth of PUM2 overexpressing plants, *PUM2* overexpression was induced by 10 μM Dex, as described previously (Aoyama and Chua, 1997; Moore et al., 2006). T3 Seeds that express either *Venus* or *PUM2-Venus* under Dex induction were surface-sterilized, resuspended in 0.1% Agar, and spotted on the surface of square MS-agar plates with 4-5 mm space in between. A total number of 64 plates (32 supplemented with 10 μM Dex by addition of 1/3000 volume of 30 mM Dex in 99.9% ethanol to the liquid MS-agar medium; 32 supplemented with 0.03% ethanol as mock control) with 16 seeds each were made, and each plate contained seeds of all the four genotypes, spotted in different order to eliminate the influence of seedling position. Plates were cold-stratified for 2 days, and vertically incubated with the seeds close to the top edge, so that the seedlings grew on the surface of the medium. Characterization of root growth was performed as described (Arribas-Hernandez et al., 2020). On DAG 2 (48h after germination), the positions of seeds and root tips were marked on the bottom side of the plates with colored markers. Positions of root tips were marked every 24 hr until DAG 11, and images of the plates were taken from the back side with a Nikon D5600 camera to show the root growth marks. The raw images were opened in Fiji and scaled. For each root, the coordinates of root growth marks were obtained by clicking on the mark using the ‘point’ tool and exported as Excel sheet. Information of individual plants (genotype and ID) were manually added to the sheet, and statistical analysis was performed in R. The coordinates were aligned relative to the DAG 0 mark (seed position) which became (0,0). Different growth indices were calculated as below, where *X* and *Y* are coordinates, and subscripts represent the DAG value:

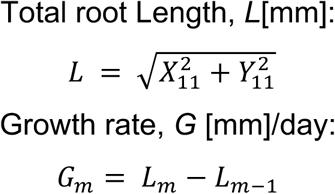

To assess the statistical significance of the observed growth indices among different genotypes, *t*-tests with Benjamini-Hochberg adjustment for multiple comparisons were performed.

All measurements can be found in Supplementary Table S4.

### Alexander staining

Alexander staining of Arabidopsis anthers was conducted as described (Alexander, 1969).The Alexander staining solution contains 10% ethanol, 25% glycerol, 0.005% (w/v) malachite green, 0.01% (w/v) acid fuchsin, 0.05% (w/v) orange G, 4% glacial acetic acid, and 1% (w/v) phenol. The staining solution was prepared in fume hood and stored in a light-protected bottle at 4°C.

2-3 Arabidopsis flowers at stage 12 (where the petals have reached a length similar to that of carpels) were selected from each genotype and dissected under a stereomicroscope on a glass slide. Only anthers were kept and the residual parts of flowers were discarded. 1 drop (∼15 µL) of Alexander staining solution was added to submerge the anthers. The sample was then covered with a cover slip and sealed with glisseal N (Borer). The slides were incubated at 50°C in a sealed, lightproof plastic box for ∼4 hr, and inspected under the stereomicroscope. Pictures were taken with 2x and 12x objectives with a SONY α6000 camera.

### PUM target identification by HyperTRIBE

The HyperTRIBE experiments were performed as described previously (Arribas-Hernández et al., 2021b). Separate HyperTRIBE experiments were performed with *PUM2*, *PUM5* and *PUM6*. For each experiment, the plant material was grown in parallel, and tissue harvest, RNA extraction, library preparation and Next-generation sequencing were performed at the same time.

DAG 8 T2 plants expressing PUM-FLAG-ADAR or free FLAG-ADAR in Col-0 wild type background were dissected and shoot apices (for PUM2 and PUM6 HyperTRIBE) or ∼3 mm long root tips (for PUM5 HyperTRIBE) were harvested in liquid nitrogen. For each genotype, 5 independent lines with similar transgene expression levels were selected for HyperTRIBE. Total RNA extraction was conducted as described above, and the RNA integrity and concentration were assessed by running Bioanalyzer RNA 6000Nano chips (Agilent).

3 µg of total RNA were shipped in dry ice to the Novogene Sequencing Center in Cambridge, UK, for library construction and sequencing (Illumina PE150, Q30≥80%). The PUM2 HyperTRIBE libraries were sequenced to a depth of at least 40 M raw reads (∼12 GB raw data), while the PUM5 and PUM6 HyperTRIBE libraries were sequenced to a depth of at least 60 M raw reads (∼18 GB raw data).

Salmon quantification (Patro et al., 2017) was conducted with the raw fastq files. The TAIR 10 Arabidopsis genome annotation (Lamesch et al., 2011), manually supplemented with the *PUM-FLAG-ADAR* and *FLAG-ADAR* sequences, was used for quasi-mapping with Salmon. In parallel, raw reads were trimmed with trimmomatic (Bolger et al., 2014) and aligned with STAR (Dobin et al., 2013). Base count files were generated with the Samtool mpileup method (Li et al., 2009), and the subsequent HyperTRIBE analysis was conducted in R with the hyperTRIBE**R** package (Rennie et al., 2021). The counts of reads containing at least one mismatch to the reference genome as well as counts of reads with no mismatches per nucleotide position were retrieved across samples, and filtered to retain high-confidence mismatch sites, defined as positions with at least 2 mismatching reads in at least 3 out of 5 PUM-FLAG-ADAR-expressing samples. The editing proportion was defined as *G*/*A*+*G*, where *A* and *G* represent the counts of reads with adenosine or guanine at the editing site, respectively. The filtering steps used to call significant A-to-G editing sites and, consequently PUM HyperTRIBE targets, were performed as described (Reichel et al., 2024), i.e. the following conditions had to be met: (1) editing proportion > 0.01 (edit was relatively frequent); 2) log2 fold change > 1 (edit was more frequent in *PUM-FLAG-ADAR* samples than in *FLAG-ADAR* samples); 3) adjusted p-value > 0.01 (editing proportion difference was significant). Specifically, for PUM2 HyperTRIBE, genes with multiple lower-confidence edit sites with an adjusted p-value between 0.01 and 0.1 or an edit proportion < 0.01 were also included in the target set. The generated target sets were then used for downstream analysis. Gene ontology enrichment analysis was conducted based on previously published functional annotations (Thomas et al., 2022). *De novo* motif discovery in the 5’ and 3’-UTR of PUM target genes were performed with HOMER (Heinz et al., 2010).

### Venn diagrams and significance of gene set overlaps

Venn diagrams were generated using the R-package eulerr (Larsson, 2018). To evaluate whether the observed overlap among gene sets was greater than expected by chance, Chi-square tests for independence were performed. Specifically, for three-way Venn diagrams, Chi-square tests for mutual independence were performed. For all statistical tests of gene set overlaps, the background set was defined as all expressed genes (tpm > 0 in any of the RNA-seq samples) in corresponding dataset.

### Selection of m^6^A-SAC-seq sites and background As

Processed data for m^6^A-SAC-seq was downloaded from (Wang et al., 2024) and 3’-UTR sites with at least 10% stoichiometry identified in seedlings were selected for further analysis. Each site was paired with a random adenosine within the 3’-UTRs of the same transcript set, ensuring a similar location distribution (that is, similar proportions through the 3’-UTR as the m^6^A sites).

### m^6^A-SAC-seq overlaps with motif sets

Counts of exact matches of each of the candidate m^6^A consensus motifs (Wei et al., 2018; Hu et al., 2021; Mao et al., 2022; Bian et al., 2024; Chen et al., 2024; He et al., 2024) were recorded using the function scan_sequences from the R package universalmotif (Tremblay, 2024), shifting each motif according to the A within the motif assumed to be the m^6^A site. As a control, this analysis was repeated similarly for the set of random adenosines (defined according to the approach stated above).

### Enrichment of motifs around m^6^A sites in PUM bound ECT2/3 targets

Stringent ECT2/3 target sets (Arribas-Hernández et al., 2021b) were split according to PUM target status based on HyperTRIBE-defined PUM2/5/6 target sets from this study. Flanking windows were selected with a width of 50 nt, excluding the 5 nt up and downstream of the site itself. For each site (m^6^A or background random A), the presence or absence of a given motif or motif set was recorded, and a generalized linear model was fit assuming a binomial response using the R package lme4 (Bates et al., 2015). The covariates were the A status (m^6^A or A) and PUM status (bound or unbound) and the interaction between these two terms. To account for the potential for correlations due to multiple sites on the same 3’-UTR, a random effect term for the gene was also included in the statistical model.

### Immunoprecipitation and mass spectrometry (IP-MS)

The *PUM2-HisLinker-Venus* and *PUM5-HisLinker-Venus* constructs were expressed in *pum2-1* and *pum5-5* mutant lines respectively, and *HisLinker-mVENUS* was expressed in Col-0 background. For each genotype, 3 independent transgenic lines were selected for IP-MS.

All procedures for immunoprecipitation were conducted in 4°C cold room and/or on ice unless specified otherwise. DAG 6 T3 whole seedlings from each line were collected and ground in liquid nitrogen. To lyse the tissue, IP buffer (50 mM Tris–HCl pH 7.5, 150 mM NaCl, 10% glycerol, 5 mM MgCl_2_, 0.1% Nonidet P40) was prepared, cooled, and supplemented with 5 mM dithiothreitol (DTT, sigma), 2 mM 4-benzenesulfonyl fluoride hydrochloride (AEBSF, Sigma), 40 μg/ml RNase A, 1 tablet of Complete protease inhibitor (Roche) and 2% plant protease inhibitor cocktail (Sigma) right before use. For each sample, ∼2 g of tissue powder were added to 8 mL IP buffer with supplements as described above and vortexed vigorously. The lysate was centrifuged at 16,000 *g* for 10 minutes, and the supernatant was passed through 0.45 μm hydrophilic filters. 45 µL of total lysate was saved for gel electrophoresis and western blot. Meanwhile, 12.5 µL of GFP-TRAP® agarose beads (Chromotek) were washed twice with cold IP buffer, and added to the lysate carefully. The lysate-bead mixture was incubated for 1 h with slow rotation. After incubation, the beads were collected by spinning for 1 min at 1,000 g. The supernatant was removed carefully with a pipette without disturbing the beads, and 45 µL supernatant was saved for gel electrophoresis and western blot. The beads were washed 4 times (1, 5, 2×10 minutes) with cold IP buffer supplemented with 5 mM DTT, and 2 times with 1x PBS buffer for 5 min each. After the final wash, 500 µL PBS buffer was added to resuspend the beads, and 200 µL mixture was saved for gel electrophoresis and western blot. The remaining 300 µL of resuspended beads were centrifuged for 1 min at 1,000 g to collect the beads. The supernatant was discarded completely and carefully with a pipette, and the beads were kept at -80 °C until mass spectrometry was performed. To assess the quality of the IP, the saved total lysate (input), supernatant after incubation, and bead fraction were mixed with 4x LDS and heated at 70 °C for 10 min. The protein samples were run on a 4-20% Criterion TGX Precast gel (Biorad). After electrophoresis, the gels were either subject to western blot as described above, or silver stained as described below.

### Preparation of samples for liquid chromatography-mass spectrometry analysis

On-bead digestion with LysC and trypsin was performed for downstream liquid chromatography-mass spectrometry analysis. The beads were resuspended in 20 µL of lysis buffer (6 M guanidium chloride (GdCl), 10 mM Tris(2-carboxyethyl)phosphine (TCEP), 40 mM 2-chloroacetamide (CAA), 50 mM HEPES pH 8.5) and sonicated for 3 x 30 s on/30 s off cycles using BioRupter Pico. The samples were centrifuged at 5,000 g for 1 min and diluted with 2x volume (40 µL) of freshly prepared digestion buffer (10% acetonitrile (ACN), 50 mM HEPES pH 8.5) and 200 ng LysC. The reactions were mixed well by vortexing and incubated at 37°C for 4 hours with shaking at 1,500 rpm. Next, 140 µL of digestion buffer and 100 ng Trypsin were added to the reactions. Tubes were vortexed and incubated at 37°C overnight with shaking at 1,500 rpm. The following day, digested samples were mixed with an equal volume of 2% trifluoro acetic acid (TFA), vortexed and centrifuged at 2,000 g for 1 min. The supernatant was transferred to a clean tube and centrifuged again to remove all bead debris from digested proteins.

The digested protein samples were purified with C-18 stage tips. A packing syringe fitted with a blunt-end needle was used to extract two layers of C18 filter material (Empore 2215, 66883-U) and gently insert them into 200 µL tips. The StageTips were activated by sequentially washing with 30 µL methanol, 30 μl Buffer B (80% ACN, 0.1% Formic Acid (FA)), and 2× 30 μl Buffer A’ (3% ACN, 1% TFA). For every wash, StageTips were centrifuged at ∼1,000 g for 60 s. Protein samples were then loaded onto StageTip, maximum 50 µL at a time. StageTips were washed twice with 100 µL Buffer A (0.1% FA), and eluted twice with 30 µL Buffer B’ (40% ACN, 0.1% FA). The eluted protein samples were vacuumed with SpeedVac Vacuum Concentrators (Thermo Scientific) at 60 °C for 1 hr, and resuspended in 12 µL Buffer A* (2% ACN, 1% TFA) with iRT peptides. 1 µL of sample was used for measuring the peptide concentration with Nanodrop.

### Liquid chromatography-mass spectrometry analysis

Non-targeted mass spectrometry was performed on a quadrupole Orbitrap benchtop mass spectrometer (Q Exactive Classic, Thermo Scientific) equipped with an EASY-nLC 1000 nano-liquid chromatography system (Thermo Scientific). All steps were performed as described previously (Tankmar et al., 2023).

### Analysis of mass spectrometry data

Raw mass spectrometry data was processed using Proteome Discoverer (v2.4, Thermo) with label-free quantification (LFQ) in both processing and consensus stages. During data processing, oxidation of methionine, acetylation at protein N-termini, and loss of methionine were included as dynamic modifications, and cysteine carbamidomethylation was set as a static modification. Results were filtered using Percolator with 1% false discovery rate (FDR), and quantification was done with the Minora feature detector. Spectral matching was conducted with SequestHT against the TAIR10 database. In total, 289 proteins were identified across all the samples. The protein abundance matrix was then exported to R for downstream analysis.

Log10 normalization was applied to the raw protein abundances. To identify proteins with differential abundance across sample groups, the R package limma was used (Smyth, 2004). A linear model was first constructed with a design and contrast matrix to define sample contrasts. Fold changes and p-values were calculated from the linear model. The fitted model was then passed to functions ‘treat’ and ‘decideTests’ were applied to the fitted model to identify differentially abundant proteins, where p-values were adjusted for multiple testing using the Benjamini– Hochberg method (Benjamini and Hochberg, 1995). The enriched proteins were considered as PUM-associated and gene ontology enrichment analysis (Thomas et al., 2022) was performed on these proteins.

### Silver staining

To estimate the amount of protein from IP fractions, a series of bovine serum albumin (BSA) dilutions was loaded on the gel for silver staining. The gel was fixed with 30% ethanol and 10% acetic acid for 2 hours, and washed with 20% ethanol for 10 min, 10% ethanol for 10 min, and water for 2 x 20 min. Next, the gel was rinsed in sensitizing solution (1 mM Na_2_S_2_O_3_) for 1 min, washed with water for 2 x 1 min, and incubated with staining solution (0.1% (w/v) silver nitrate) for 1 hr with shaking. Then, the gel was washed with water for 2 x 1 min and developed in freshly prepared developing solution (2% (w/v) Na_2_CO_3_, 0.04% formaldehyde prepared by diluting 37% formaldehyde 1000-fold into the developing solution right before use). When the bands became clearly visible, the reaction was stopped with 2% acetic acid. The gel was transferred on to a glass plate and an image of the stained gel was taken with a Nikon D5600 camera.

### Accession numbers

mRNA sequencing data sets generated in this study were submitted to the European Nucleotide Archive (ENA) under accession number PRJEB89231 (Submission ERA33116164). IP-MS data were submitted to the EMBL-PRIDE repository **IP-MS data to EMBL-PRIDE** under project accession PXD063607.

Reviewers can access the dataset by logging in to the PRIDE website using the following account details:

Username

Password: YgXCDariqwVx

## ACKNOWLEDGEMENTS

We thank Theo Bölsterli and Rene Hvidbjerg for plant care. The Nottingham Arabidopsis Stock Centre is thanked for providing T-DNA insertion lines. Rasmus Kjøller and Rasmus Hartmann-Petersen are thanked for kind gifts of chemicals needed for Alexander staining. The DTU Proteomics Core Facility is thanked for conduction of LC-MS experiments. This work was supported by a Hallas Møller Ascending Investigator grant (NNF19OC0054973) from the Novo Nordisk Foundation to P.B., an Emerging Investigator grant from the Novo Nordisk Foundation (NNF24SA0092867) to S.R., and by an infrastructure grant (CF20-0659) from Carlsberg Fondet to P.B.

## AUTHOR CONTRIBUTIONS

J.C. carried out all experimental work and all data analyses except those reported in Figures 3C-E, M.S.A. analyzed data, S.R. supervised data analysis by J.C. and M.S.A. and carried out the motif analyses reported in Figure 3D-E, P.B. and L.A.-H. designed the study and supervised the work, P.B. wrote the paper with input from all authors.

## SUPPLEMENTARY DATA

**Supplementary Figure S1:** Expression of Arabidopsis PUM1-PUM6 from published RNA-seq data.

**Supplementary Figure S2:** PUM2/5/6-FLAG-ADAR expression in the Arabidopsis stable transgenic lines used for HyperTRIBE.

**Supplementary Figure S3:** Transgenic expression of *PUM2/5/6-FLAG-ADAR* constructs does not have major impact on plant development or overall transcriptome

**Supplementary Figure S4:** Workflow of PUM2/5/6-HyperTRIBE.

**Supplementary Figure S5:** PUM HyperTRIBE (HT) efficiently identifies edit sites with significantly differential editing proportion with minimum biases.

**Supplementary Figure S6:** PUM2/5/6 target sets overlap with m^6^A-containing transcripts and ALBA2/4 targets.

**Supplementary Figure S7:** Neither the single knockout of *PUM2/3/5/6* nor the combined knockout of *PUM2-PUM3, PUM3-PUM6* or *PUM5-PUM6* produce significant defects in early-stage leaf development.

**Supplementary Figure S8:** PUM2 overexpression represses leaf, shoot and root growth.

**Supplementary Figure S9:** CRISPR-Cas9-induced knockout of *ECT2* does not cause conspicuous growth defects.

**Supplementary Figure S10:** PUM2-mediated developmental defect is more pronounced in the triple *ect2 ect3 ect4* mutant background.

**Supplementary Figure S11:** Transcriptome-wide analysis of PUM2 binding to mRNAs in the presence or absence of ECT2/3/4.

**Supplementary Figure S12:** Identification of PUM2 and PUM5 interacting proteins by IP-MS.

**Supplementary Figure S13:** Lack of evidence for functional physical association between PUM2/5 and ECTs (extended data).

**Supplementary Table S1:** List of plants used in this study.

**Supplementary Table S2:** List of molecular cloning components (vectors, primers, constructs) used in this study.

**Supplementary Table S3:** List of software used in this study.

**Supplementary Table S4:** Growth measurement of PUM^OX^ plants.

**Supplementary Table S5:** PUM HyperTRIBE results.

**Supplementary Table S6:** PUM IP-MS results.

