## Supplementary Figure S1 for "PUMILIOs and m^6^A-ECT2/ECT3 share mRNA targets and exert opposing control over organogenesis"

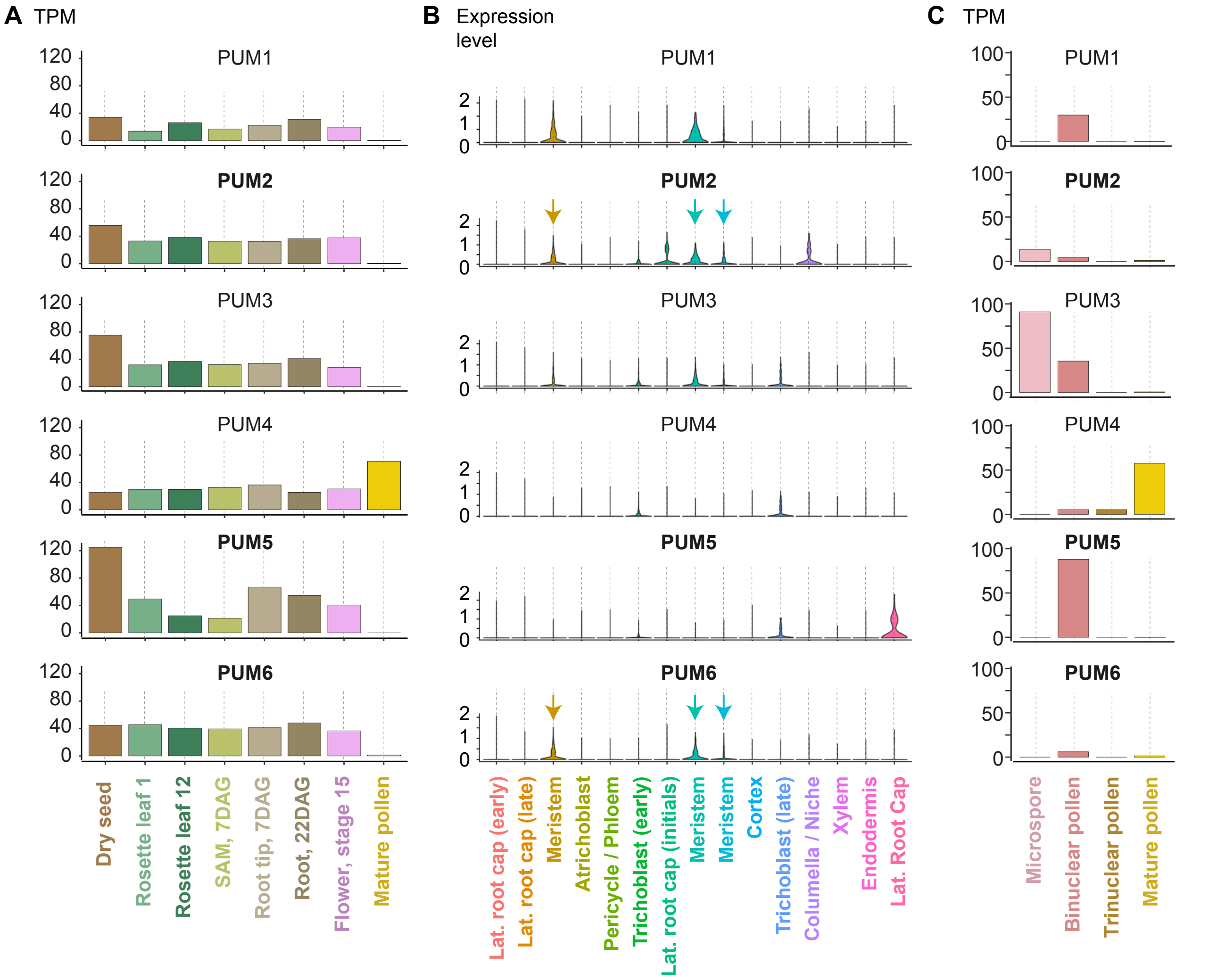


**Supplementary Figure S1. Expression of Arabidopsis PUM1-PUM6 from published RNA-seq data.** **(A)** *PUM1-6* expression in transcripts per million (TPM) in selected tissues from bulk RNA-seq data (Mergner et al., 2020). DAG, days after germination. **(B)** *PUM1-6* expression in roots from single-cell RNA-seq data (Denyer et al., 2019). Arrows highlight the expression of *PUM2* and *PUM6* in the root meristem. **(C)** *PUM1-6* expression during the different stages of pollen maturation according to single-anther RNA-seq (microspore, binuclear and trinuclear pollen) (Le Lievre et al., 2023) and bulk RNA-seq (mature pollen) (He et al., 2019).


- Denyer, T., Ma, X., Klesen, S., Scacchi, E., Nieselt, K., & Timmermans, M. C. P. (2019). Spatiotemporal Developmental Trajectories in the Arabidopsis Root Revealed Using High-Throughput Single-Cell RNA Sequencing. Developmental Cell, 48(6), 840-852.e5. https://doi.org/10.1016/j.devcel.2019.02.022
- He, S., Vickers, M., Zhang, J., & Feng, X. (2019). Natural depletion of histone H1 in sex cells causes DNA demethylation, heterochromatin decondensation and transposon activation. ELife, 8. https://doi.org/10.7554/eLife.42530
- Le Lievre, L., Chakkatu, S. P., Varghese, S., Day, R. C., Pilkington, S. M., & Brownfield, L. (2023). RNA-seq analysis of synchronized developing pollen isolated from a single anther. Frontiers in Plant Science, 14. https://doi.org/10.3389/fpls.2023.1121570
- Mergner, J., Frejno, M., List, M., Papacek, M., Chen, X., Chaudhary, A., Samaras, P., Richter, S., Shikata, H., Messerer, M., Lang, D., Altmann, S., Cyprys, P., Zolg, D. P., Mathieson, T., Bantscheff, M., Hazarika, R. R., Schmidt, T., Dawid, C., … Kuster, B. (2020). Mass-spectrometry-based draft of the Arabidopsis proteome. Nature, 579(7799), 409–414. https://doi.org/10.1038/s41586-020-2094-2
