## Supplementary Figure S2 for "PUMILIOs and m^6^A-ECT2/ECT3 share mRNA targets and exert opposing control over organogenesis"

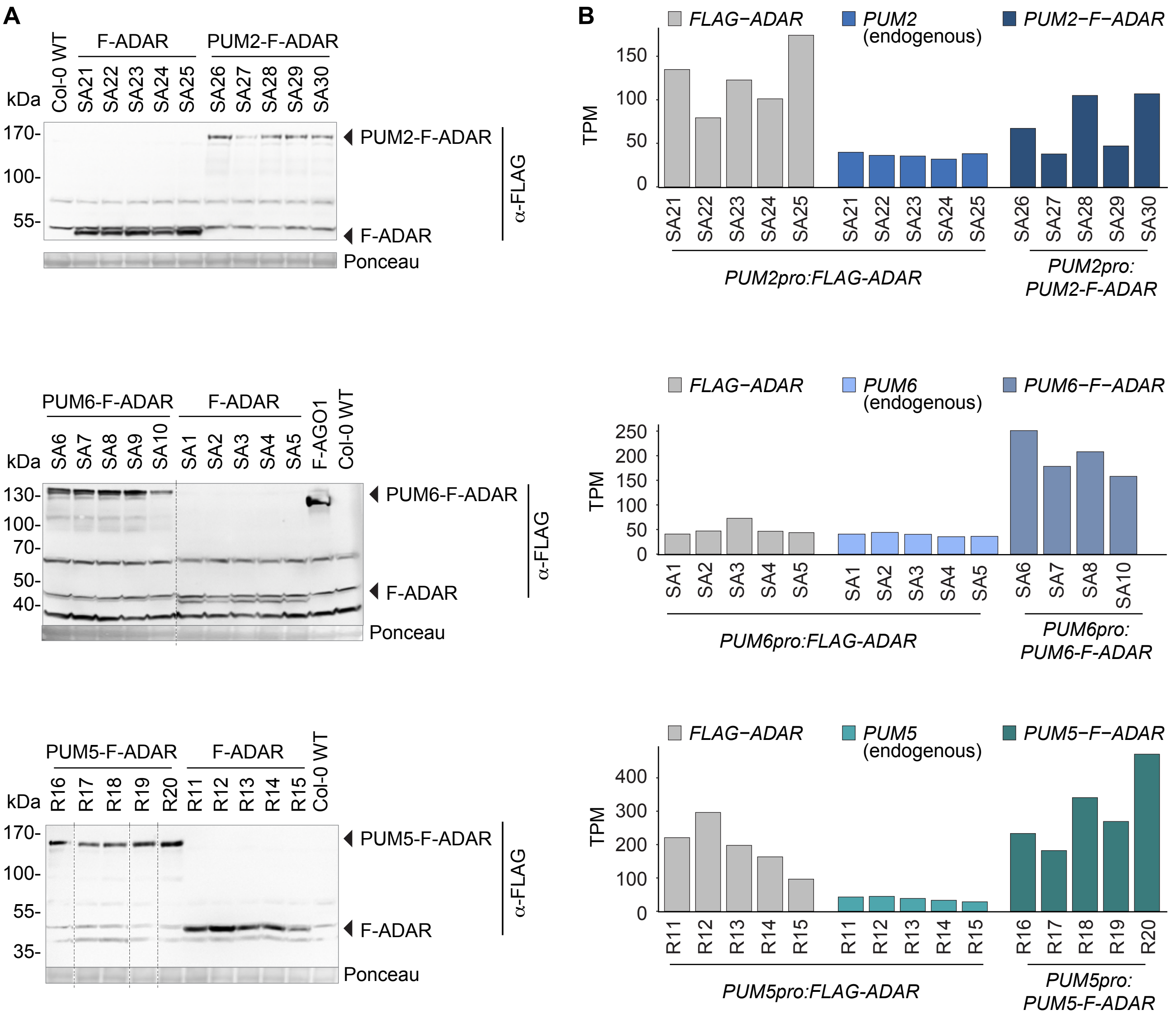


**Supplementary Figure S2. PUM2/5/6-FLAG-ADAR expression in the Arabidopsis stable transgenic lines used for HyperTRIBE.**

**(A)** Protein blot using anti-FLAG antibodies reveals the accumulation of PUM2/5/6-FLAG-ADAR and free FLAG-ADAR in aerial (PUM2, PUM6) or root (PUM5) tissues of 8-day-old seedlings of the transgenic lines used for HyperTRIBE. Genotypes used as controls, such as Col-0 WT (negative) and FLAG-AGO1 (positive) are indicated, and Ponceau staining is used as loading control. Vertical dotted lines separates lanes on the same membrane. **(B)** Transcript per million (TPM) of endogenous *PUM2/5/6*, transgenic *PUM2/5/6-FLAG-ADAR*, and *FLAG-ADAR* detected by mRNA-seq in the samples used for HyperTRIBE.
