## Supplementary Figure S3 for "PUMILIOs and m^6^A-ECT2/ECT3 share mRNA targets and exert opposing control over organogenesis"

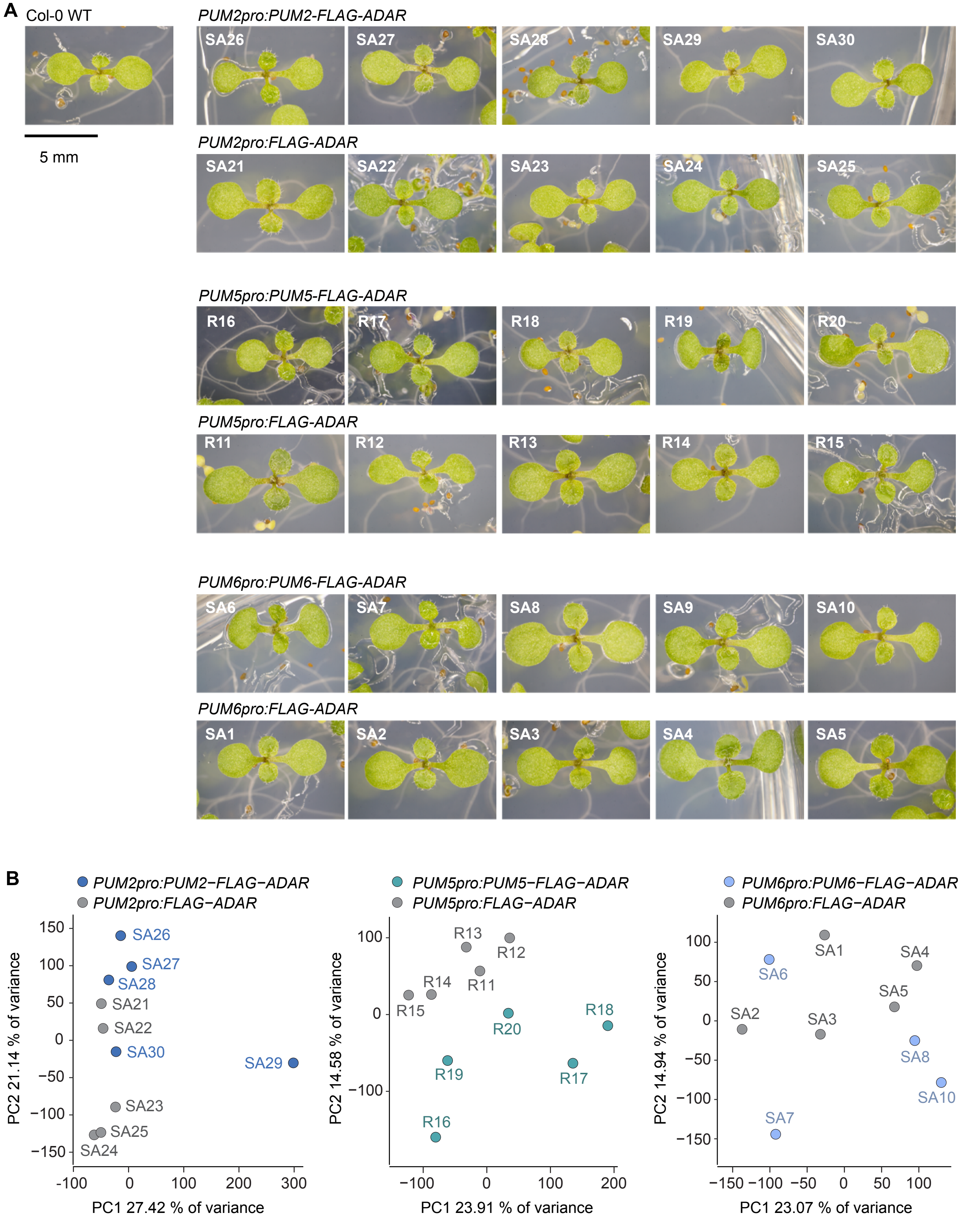


**Supplementary Figure S3. Transgenic expression of *PUM2/5/6-FLAG-ADAR* constructs does not have major impact on plant development or overall transcriptome.**
**(A)** 8-day-old seedlings of Col-0 wild type (WT) and the transgenic lines used for HyperTRIBE, expressing either PUM2/5/6-FLAG-ADAR or free FLAG-ADAR. **(B)** Principal component analysis of transcript abundance (RNA-seq) in the genotypes described in (A).
