## Supplementary Figure S4 for "PUMILIOs and m^6^A-ECT2/ECT3 share mRNA targets and exert opposing control over organogenesis"

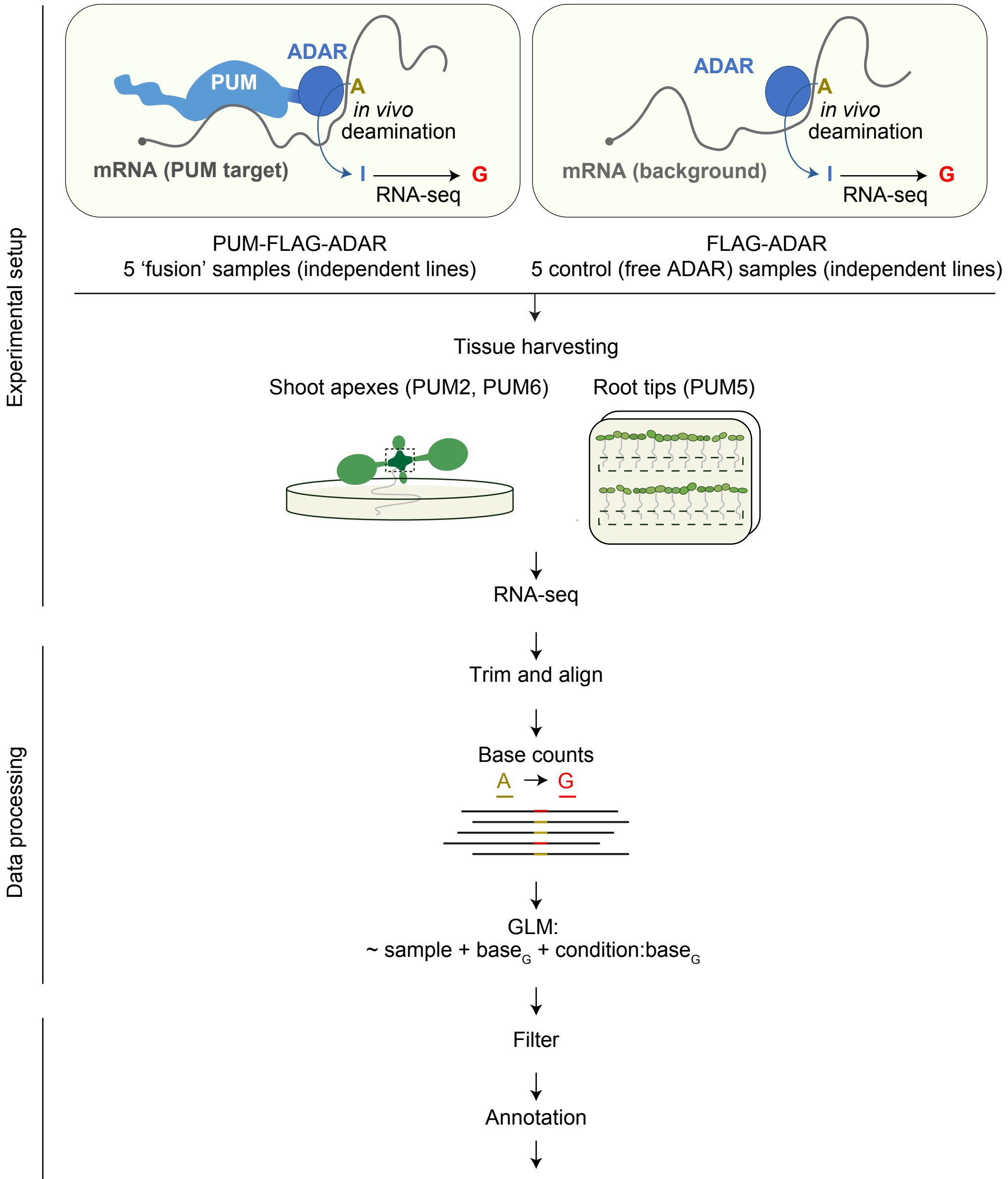


| position | gene | p_adj | edit_proportion | ... |
|---|---|---|---|---|
| 1_33941 | LHY | 2.607e-03 | 0.0290 | ... |
| 1_46478 | AT1G01080 | 8.615e-06 | 0.0906 | ... |
| 1_48692 | PDH-E1 alpha | 2.679e-05 | 0.0033 | ... |
| ... | ... | ... | ... | ... |

**Supplementary Figure S4. Workflow of PUM2/5/6-HyperTRIBE**
PUM2/5/6 fused to the catalytic domain of a hyperactive point mutant of ADAR, a deaminase from Drosophila melanogaster that causes adenine-to-inosine (A-I) editing in mRNA, are expected to edit adenosines adjacent to PUM2/5/6 binding sites *in vivo* when the FLAG-tagged fusion proteins are expressed in stable Arabidopsis transgenic lines. Editing sites are detected as adenine-to-guanine (A-G) mutations by RNA-seq, and transcripts with significantly higher PUM2/5/6-FLAG-ADAR-dependent A-to-G editing proportions compared to those caused by random editing by free FLAG-ADAR (background control) were considered PUM2/5/6 targets. For each genotype, 5 independent transgenic lines were used as biological replicates. GLM, generalized linear model.
