## Supplementary Figure S5 for "PUMILIOs and m^6^A-ECT2/ECT3 share mRNA targets and exert opposing control over organogenesis"

**A**

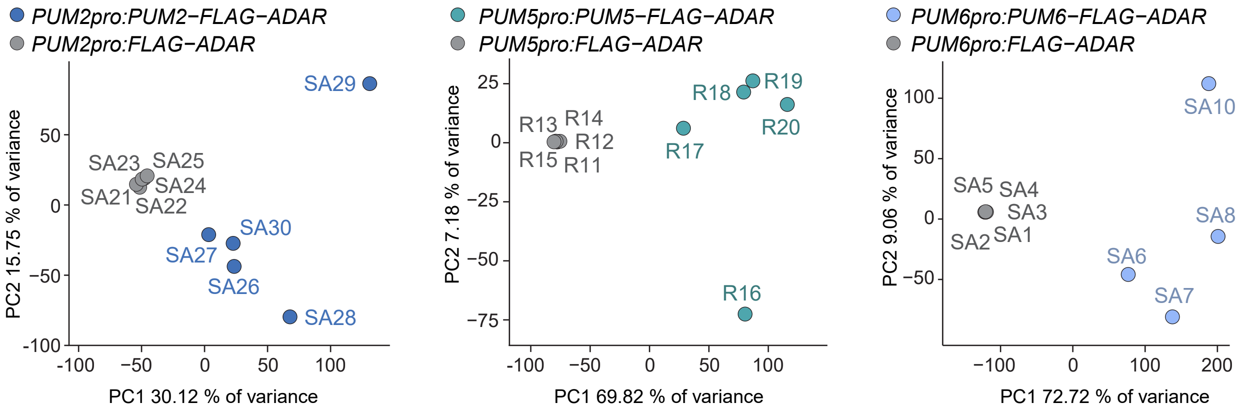


**B**

| Gene | Gene | PUM2 HT (aerial tissues) | PUM5 HT (root) | PUM6 HT (aerial tissues) |
|---|---|---|---|---|
| AT5G24490 | 30S ribosomal protein | $5.14 \times 10^{-35}$ | $1.75 \times 10^{-19}$ | $5.77 \times 10^{-89}$ |
| AT3G47470 | CAB4 | $2 \times 10^{-7}$ | $5.24 \times 10^{-8}$ | $3.11 \times 10^{-21}$ |
| AT4G36040 | DJC23 | $3.8 \times 10^{-5}$ | Non-target | Non-target |
| AT1G75820 | CLV1 | Non-target | $9.65 \times 10^{-4}$ | $9.26 \times 10^{-35}$ |
| AT3G63500 | OBE4 | Non-target | Non-target | $7.65 \times 10^{-9}$ |
| AT5G43810 | ZLL/AGO10 | Non-target | Non-target | Non-target |
| AT2G17950 | WUS/PGA6 | Non-target | Non-target | Non-target |
| AT5G64630 | FAS2 | Non-target | Non-target | Non-target |
| AT4G39090 | RD19 | Non-target | Non-target | Non-target |

**Supplementary Figure S5. PUM HyperTRIBE (HT) efficiently identifies edit sites with significantly differential editing proportion with minimum biases.**
**(A)** Principal component analysis of editing proportion per site across PUM2/5/6 HyperTRIBE samples. **(B)** Previously reported PUM2 targets identified by yeast 3-hybrid screening (Francischini and Quaggio, 2009). Numbers are adjusted p-values of their confidence as PUM targets according to HyperTRIBE (this work).
