## Supplementary Figure S6 for "PUMILIOs and m^6^A-ECT2/ECT3 share mRNA targets and exert opposing control over organogenesis"

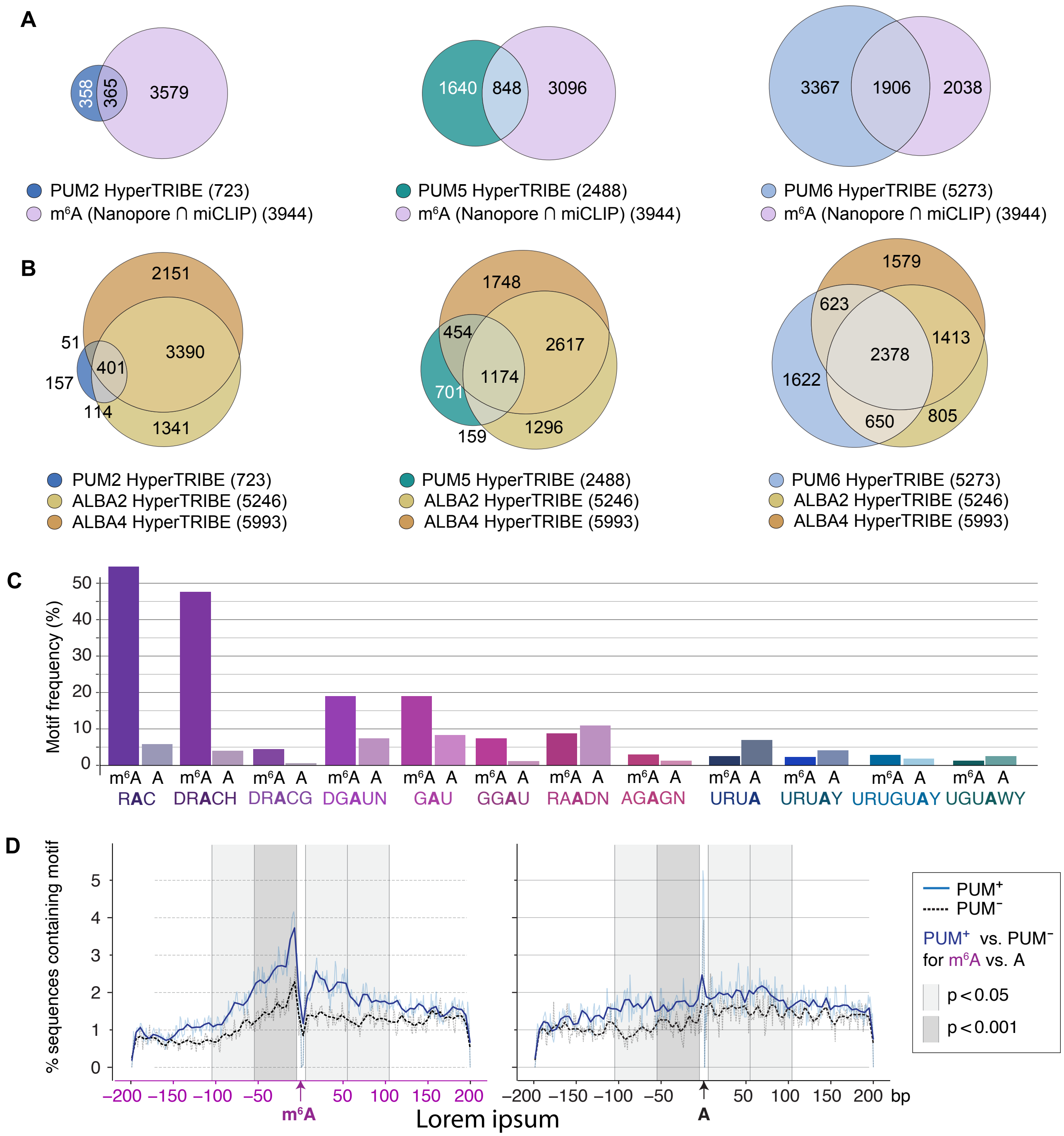


**Supplementary Figure S6. PUM2/5/6 target sets overlap with m$^6$A-containing transcripts and ALBA2/4 targets**
**(A-B)** Overlap between PUM2/5/6 targets identified by HyperTRIBE and (A) m$^6$A-containing transcripts identified by miCLIP and Nanopore (Parker et al., 2020) or (B) ALBA2 and ALBA4 targets determined by HyperTRIBE and TRIBE respectively (Reichel et al., 2024). **(C)** Percentage of m$^6$A sites in 3'-UTRs contained in various motifs previously found to be enriched around m$^6$A as described in Fig. 3C. **(D)** Percentage of seedling-derived 3'-UTR m$^6$A-SAC-seq sites (Wang et al., 2024) (left panel) or random As in ECT2/3 target mRNAs sampled from similar locations (right panel) with PRE-like motifs (URUAY, UGUAWY, UGUAH, UGUA, URUGUAY) present at given distances from the site, split by whether the transcript is PUM2/5/6 HT target (PUM$^+$) or not (PUM$^-$). Dark lines are smooth representations of the raw data (dim lines). Bands in different shades of grey represent windows of 50-nt width around the site (excluding the immediately adjacent 5 nt).The associated p-values indicate the significance of the difference in PRE-like motif depletion in the PUM$^+$ to PUM$^-$ subset comparisons with sequence windows centered either on verified m$^6$A sites or random As.


- Parker, M.T., Knop, K., Sherwood, A.V., Schurch, N.J., Mackinnon, K., Gould, P.D., Hall, A.J.W., Barton, G.J., and Simpson, G.G. (2020). Nanopore direct RNA sequencing maps the complexity of Arabidopsis mRNA processing and m6A modification. eLife 9, e49658.
- Reichel, M., Tankmar, M.D., Rennie, S., Arribas-Hernández, L., Lewinski, M., Köster, T., Wang, N., Millar, A.A., Staiger, D., and Brodersen, P. (2024). ALBA proteins facilitate cytoplasmic YTHDF-mediated reading of m6A in Arabidopsis. EMBO J., 1-30-30.
- Wang, G., Li, H., Ye, C., He, K., Liu, S., Jiang, B., Ge, R., Gao, B., Wei, J., Zhao, Y., Li, A., Zhang, D., Zhang, J., and He, C. (2024). Quantitative profiling of m6A at single base resolution across the life cycle of rice and Arabidopsis. Nat. Commun. 15, 4881.
