## Supplementary Figure S7 for "PUMILIOs and m^6^A-ECT2/ECT3 share mRNA targets and exert opposing control over organogenesis"

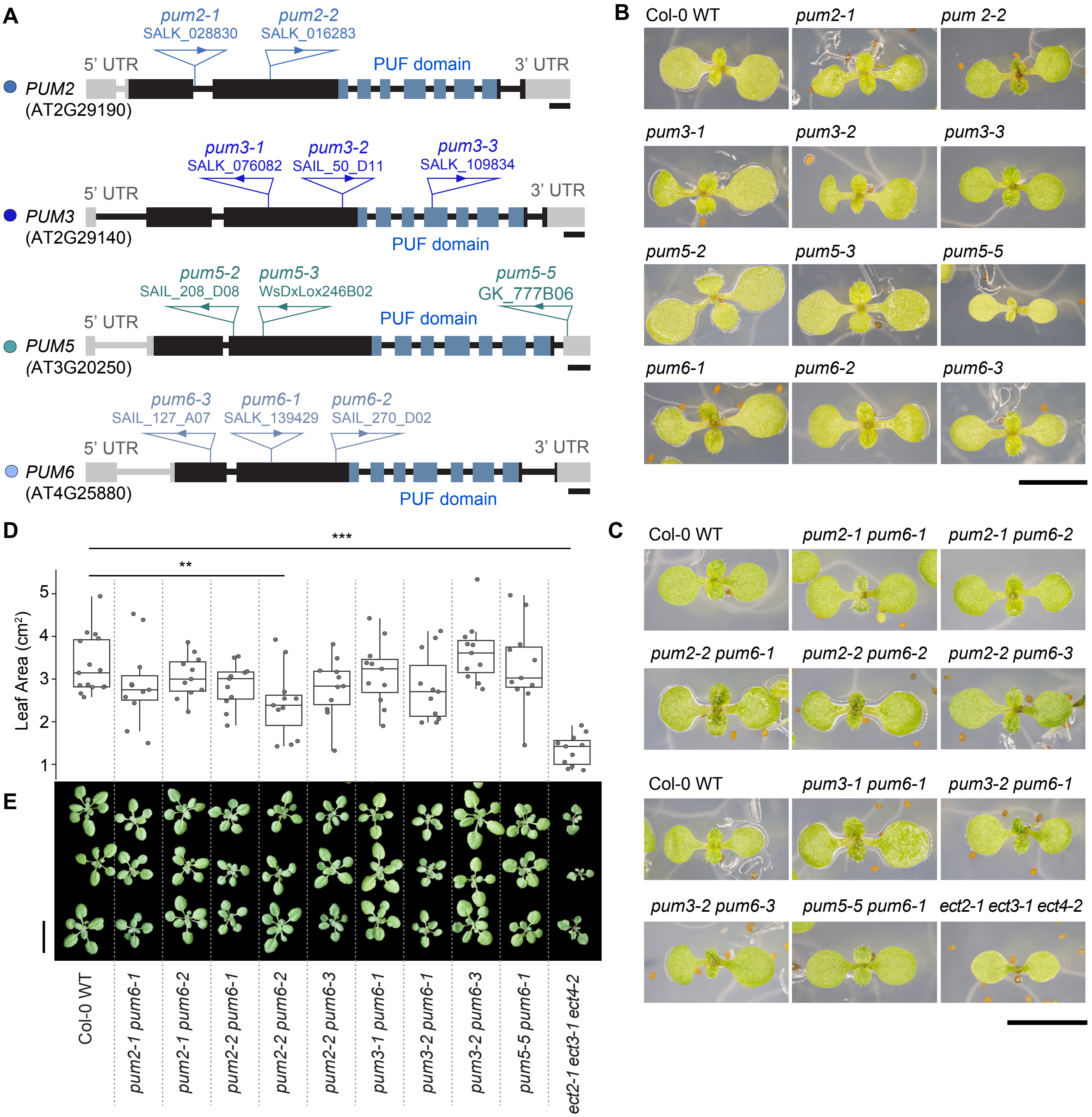


**Supplementary Figure S7. Neither the single knockout of PUM2/3/5/6 nor the combined knockout of PUM2-PUM3, PUM3-PUM6 or PUM5-PUM6 produce significant defects in early-stage leaf development.** **(A)** Gene models of *PUM2/3/5/6* marking the position of the T-DNA insertions of the indicated alleles. Scale bars, 200 bp. **(B-C)** 1-week-old seedlings of *pum2/3/5/6* single mutants (B), and *pum2 pum3, pum3 pum6* and *pum5 pum6* double mutants (C). Col-0 wild type (WT) and *ect2 ect3-ect4* genotypes are included as reference for the timing of emergence of the first pair of true leaves. Scale bars, 4 mm. **(D-E)** Representative rosettes of 2.5-week-old *pum* double mutants (D) and quantification of their total leaf area. (E). ~10 plants from each line were examined. **, p-adj < 0.01; ***, p-adj < 0.001 for Wald t-tests with Benjamini-Hochberg correction for multiple comparisons. Scale bar, 2 cm.
