## Supplementary Figure S8 for "PUMILIOs and m^6^A-ECT2/ECT3 share mRNA targets and exert opposing control over organogenesis"

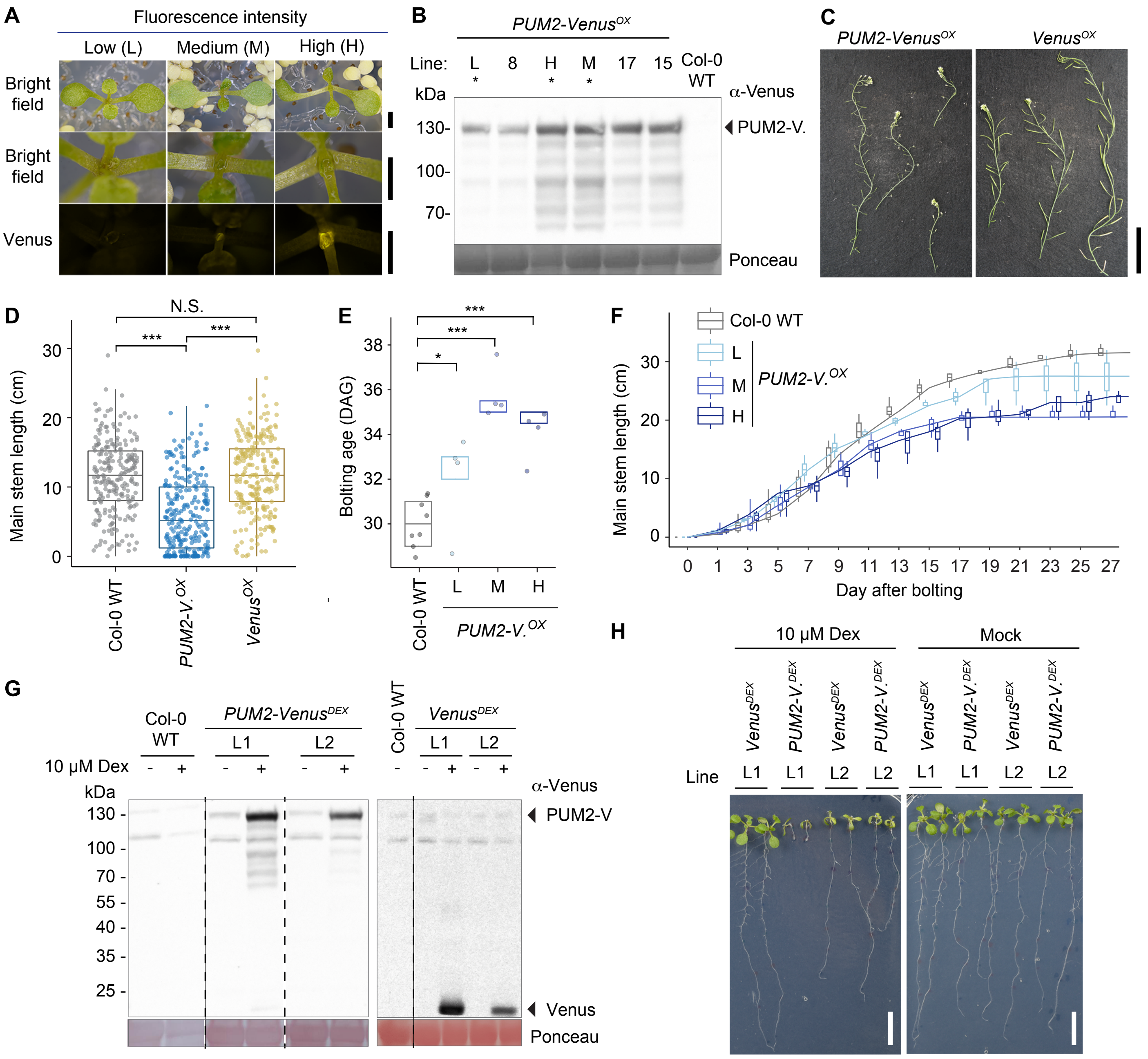


**Supplementary Figure S8. PUM2 overexpression represses leaf, shoot and root growth**

**(A)** Fluorescence microscopy of 8-day-old T1 seedlings overexpressing (ox) PUM2-Venus (*Pro*$_{uS7\gamma}$:*PUM2-Venus*) at different levels. Scale bars, 1 mm. **(B)** Protein blot analysis of T2 seedlings of several independent *PUM2-Venus*$^{OX}$ stable transgenic lines. Lines with relatively low (L), medium (M) and high (H) expression, marked with an asterisk, were used for subsequent analyses. Ponceau staining is used as loading control. **(C-D)** Characterization of main stem growth in T1 plants overexpressing either PUM2-Venus or free Venus in the Col-0 ecotype. (C) Main stem of 7-week-old plants. Scale bar, 5 cm. (D) Quantification of main stem length in 5-week-old plants. Around 380 plants per genotype were examined. Wild type Col-0 plants were included as reference. N.S., not significant; ***, p-adj < 0.001 for t-test with Benjamini-Hochberg corrections for multiple comparisons. **(E-F)** Characterization of the reproductive phenotype exhibited by T2 plants of the L, M, and H *PUM2-Venus*$^{OX}$ lines selected in (B). (E) Bolting age in days after germination (DAG). *, p-adj < 0.05; ***, p-adj < 0.001 for t-tests with Benjamini-Hochberg corrections. (F) Quantification of main stem length after bolting over time. **(G-H)** Characterization of 11-day-old T2 seedlings of two independent dexamethasone (Dex)-inducible *PUM2-Venus*$^{DEX}$ and *Venus*$^{DEX}$ lines (Fig. 4A) with or without 10 μM Dex in the growth media. Equal volume of the dexamethasone solvent (ethanol) was used for the mock (Dex -) samples. (G) Protein blot analysis. Ponceau staining was used as loading control. (H) Root phenotype. Scale bars, 1 cm.
