## Supplementary Figure S9 for "PUMILIOs and m^6^A-ECT2/ECT3 share mRNA targets and exert opposing control over organogenesis"

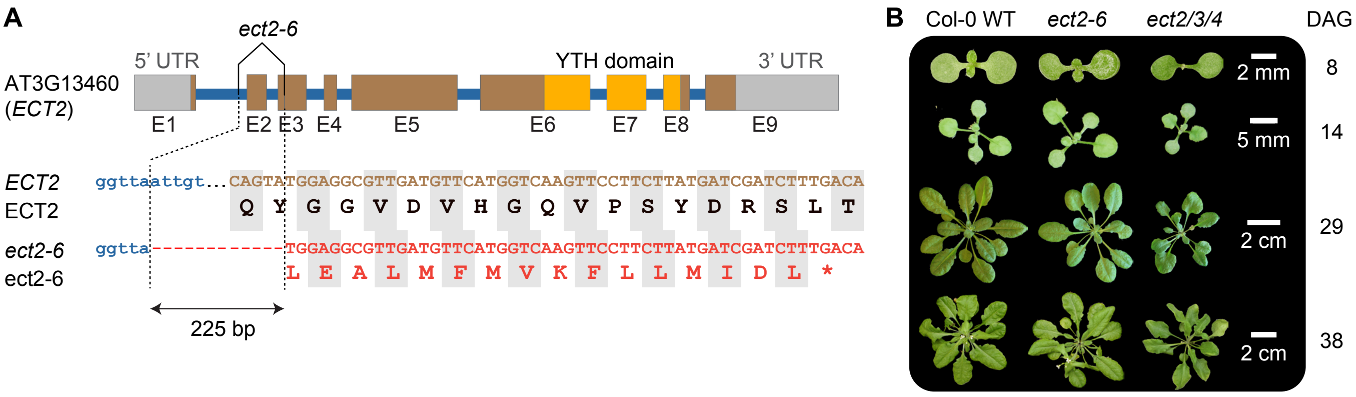


**Supplementary Figure S9. CRISPR-Cas9-induced knockout of ECT2 does not cause conspicuous growth defects**
**(A)** Schematic representation of *ect2-6*, an *ECT2* mutant allele produced by CRISPR-Cas9. A 225 bp-deletion in *ECT2* causes the absence of exon 2 (E2) and a frameshift that results in a premature stop codon in exon 3 (E3). **(B)** Photographs of *ect2-6* plants at different growth stages. Col-0 wild type and the *ect2-3/ect3-2/ect4-2* (*ect2/3/4*) triple mutant plants are included as controls. DAG, days after germination.
