## Supplementary Figure S10 for "PUMILIOs and m^6^A-ECT2/ECT3 share mRNA targets and exert opposing control over organogenesis"

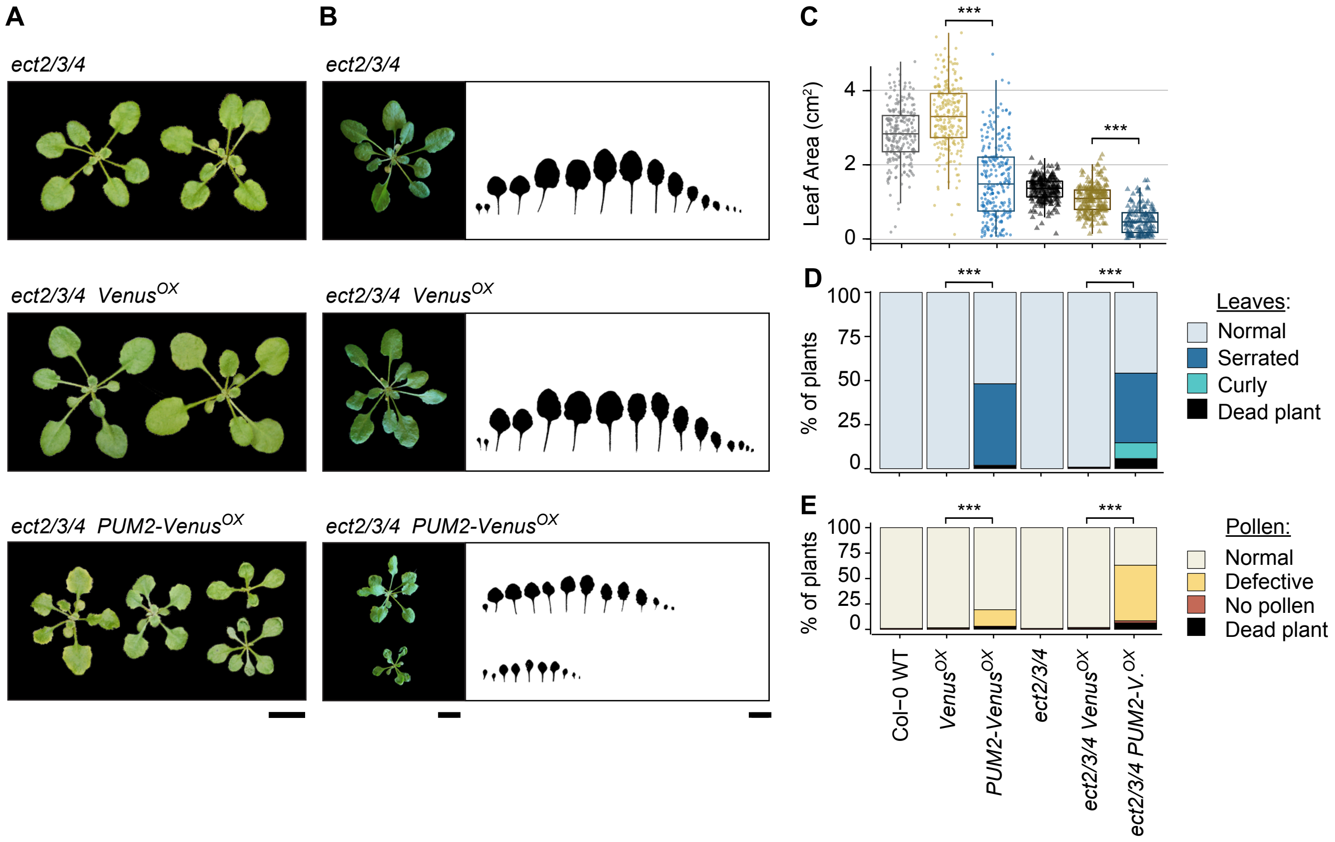


**Supplementary Figure S10. PUM2-mediated developmental defect is more pronounced in the triple *ect2 ect3 ect4* mutant background.**

**(A)** Rosettes of representative 3.5-week-old Arabidopsis plants of Col-0 wild type (WT), the triple mutant *ect2-3 ect3-2 ect4-2* (*ect2/3/4*) (Arribas-Hernández et al., 2020), and and transgenic lines (T1 plants) expressing PUM2-Venus$^{OX}$ or Venus$^{OX}$ (Fig. 4A, top panel) in these two genetic backgrounds. **(B)** Leaf profiling of 4-week-old plants of the same genotypes as in (A). (**C**) Quantification of rosette size of T1 plants of the genotypes in (A,B). Statistical significance was assessed using Student's t-test with Benjamini-Hochberg corrections. **(D,E)** Proportions of T1 plants described in (A-C) falling in different categories according to leaf morphology (D) or pollen appearance (E). For D, E: Statistical significance was assessed using Chi-square test with Benjamini-Hochberg corrections. Asterisks for significance for C-E: ***, p-adj < 0.001. Scale bars, 1 cm.

Arribas-Hernández, L., Simonini, S., Hansen, M.H., Paredes, E.B., Bressendorff, S., Dong, Y., Østergaard, L., and Brodersen, P. (2020). Recurrent requirement for the m$^6$A-ECT2/ECT3/ECT4 axis in the control of cell proliferation during plant organogenesis. Development 147, dev189134.
