## Supplementary Figure S11 for "PUMILIOs and m^6^A-ECT2/ECT3 share mRNA targets and exert opposing control over organogenesis"

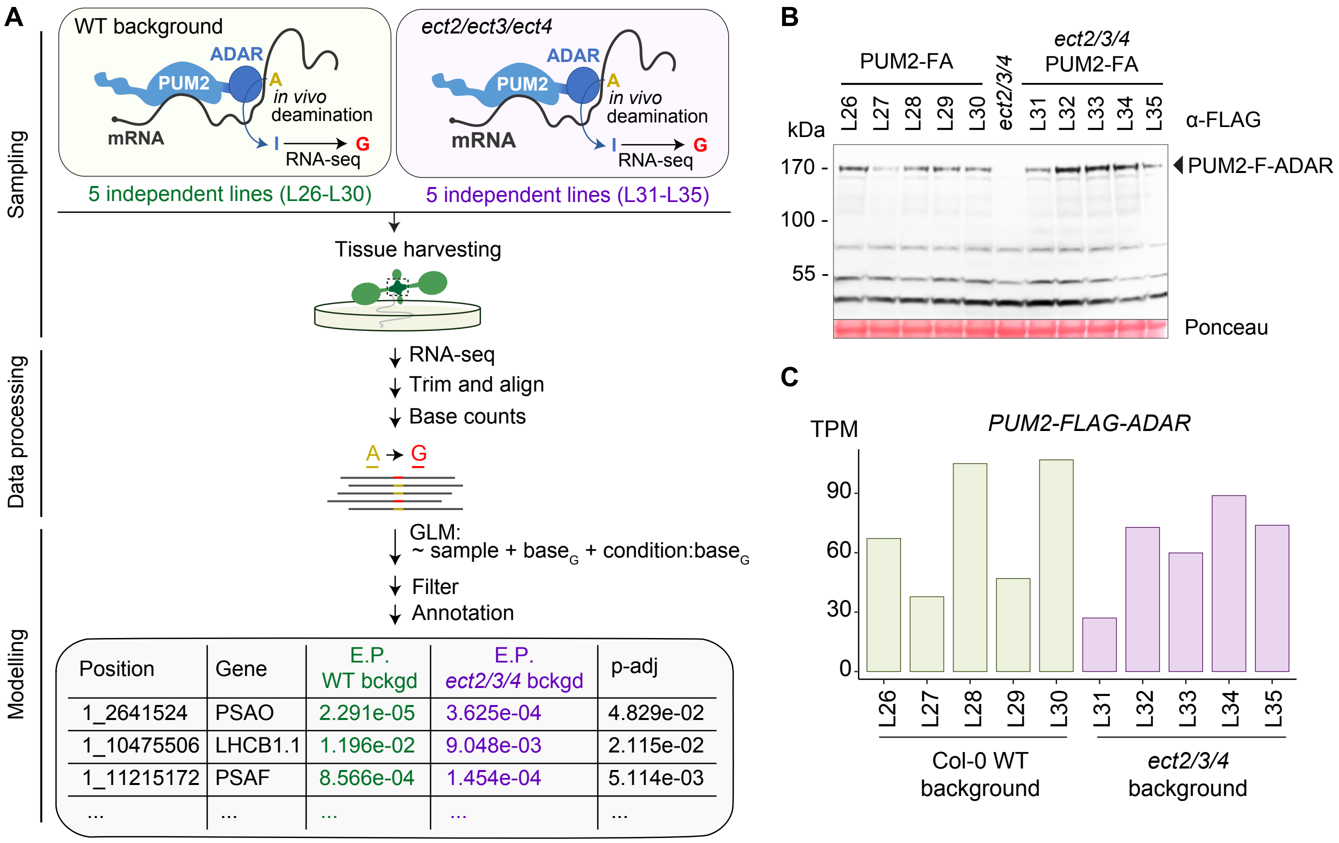


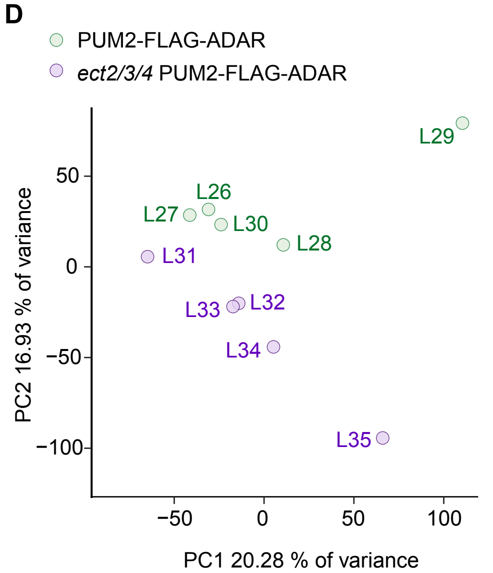


**E**

| | AGI | Gene | Edited site | p-adj | E.P. Fold Change |
|---|---|---|---|---|---|
| Photosynthesis | AT1G31330 | PSAF | 1_11215172 | 5.11E-03 | 2.53 |
| | AT1G61520 | LHCA3 | 1_22700873 | 6.02E-02 | 1.61 |
| | AT1G77590 | LACS9 | 1_29149594 | 5.88E-02 | 0.74 |
| | AT1G08380 | PSAO | 1_2641524 | 4.83E-02 | -3.94 |
| | AT1G29930 | LHCB1.3 | 1_10478390; 1_10478425 | 1.99E-02; 9.66E-02 | -2.68; -3.31 |
| | AT4G10000 | AT4G10000 | 4_6260692 | 7.89E-06 | -2.88 |
| | AT1G80920 | DJC22 | 1_30403787 | 4.46E-02 | -2.70 |
| Ribosome | AT3G11510 | uS11y | 3_3623984 | 9.66E-02 | 3.19 |
| | AT5G30510 | bS1c | 5_11620241 | 5.10E-10 | -1.87 |
| Transport | AT3G14120 | NUP107 | 3_4684766 | 3.20E-02 | 1.40 |
| | AT2G33330 | PDLP3 | 2_14123109 | 6.72E-02 | -3.77 |
| Cytoskeleton | AT1G06060 | AT1G06060 | 1_1834616 | 6.81E-02 | -1.81 |
| Stress response | AT5G42520 | BPC6 | 5_17002060 | 2.10E-03 | 1.70 |
| | AT3G05727 | DEG28 | 3_1693852 | 4.42E-02 | 1.68 |
| Chromosome | AT4G15890 | CAP-D3 | 4_9022744 | 9.43E-02 | -1.64 |
| Other | *AT3G16410* | *NSP4* | *3_5573669* | *8.72E-02* | *2.68* |
| | AT5G16660 | AT5G16660 | 5_5465643 | 3.74E-02 | 1.25 |

**Supplementary Figure S11. Transcriptome-wide analysis of PUM2 binding to mRNAs in the presence or absence of ECT2/3/4**

**(A)** Experimental design to identify differences in PUM2 binding to target mRNAs between wild type (WT) and triple *ect2/ect3/ect4* mutant plants by comparative HyperTRIBE (HT). GLM, generalized linear model. E.P., editing proportion. **(B,C)** Characterization of PUM2-FLAG-ADAR (PUM2-FA) expression levels in the transgenic lines (L26-L35) used for the comparative HT described in (A), by protein blot analysis (B) and RNA-seq (C). **(D)** Principal component analysis of editing proportions in the comparative HT experiment described in (A). **(E)** List of genes with significantly different editing proportions (E.P.), at the indicated sites, by PUM2-FLAG-ADAR in an *ect2/ect3/ect4* background compared to a wild type one. The genes are grouped by the indicated functional categories.
