## Supplementary Figure S12 for "PUMILIOs and m^6^A-ECT2/ECT3 share mRNA targets and exert opposing control over organogenesis"

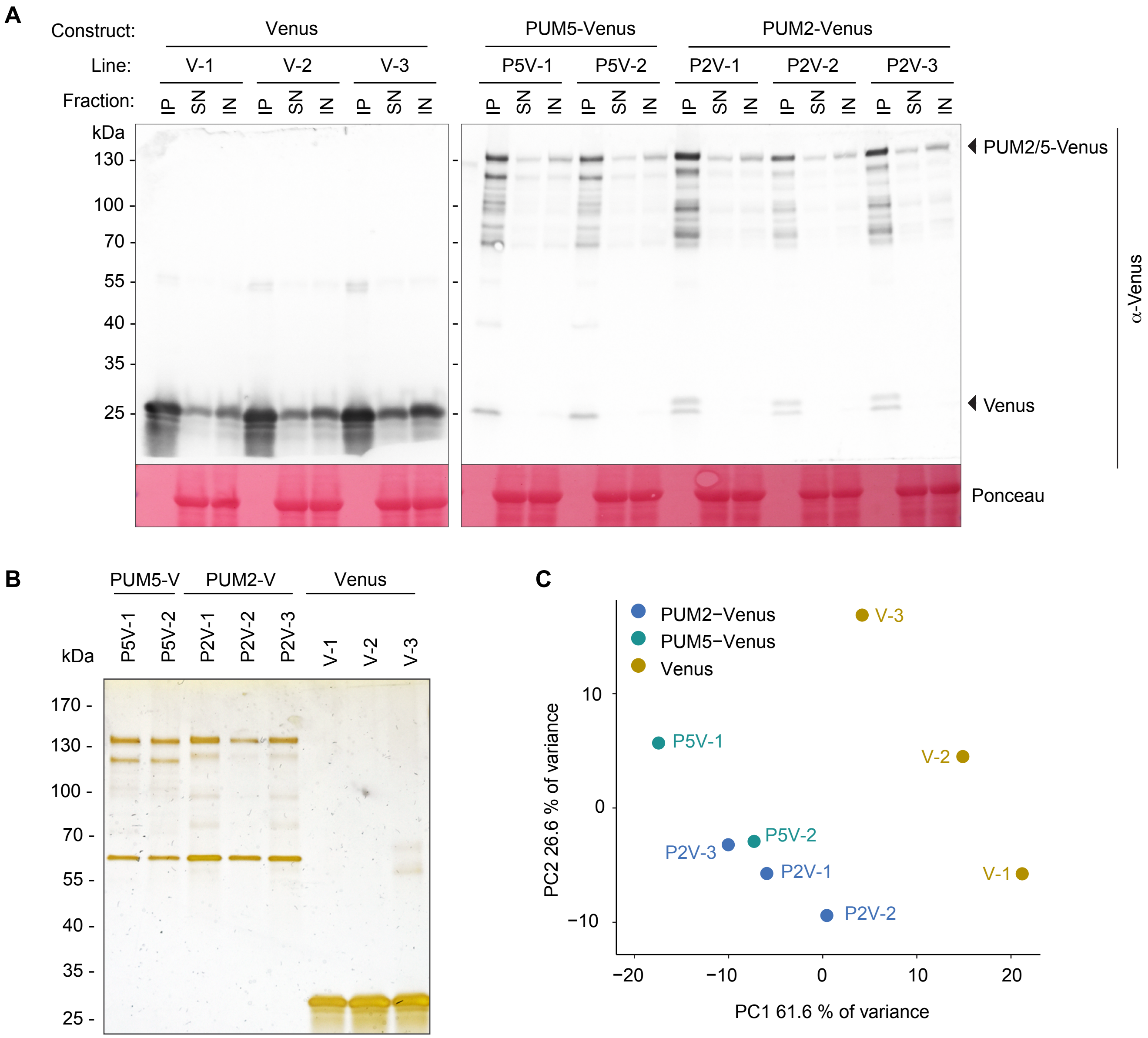


**Supplementary Figure S12. Identification of PUM2 and PUM5 interacting proteins by IP-MS**
**(A)** Protein blot analysis of GFP-Trap immunopurified (IP), supernatant (SN) and input (total lysate) fractions from total lysates of 8 day-old whole seedlings of independent transgenic lines expressing free Venus (V-1 to -3), PUM5-Venus (P5V-1 and -2) or PUM2-Venus (P2V-1 to -3). Ponceau staining of the membrane is used as loading control for the input and supernatant samples. **(B)** Silver staining of the IP fractions in (A). **(C)** Principal component analysis of peptide abundance identified by mass spectrometry in the different Venus and PUM2/5-Venus IP samples shown in (B).
