## Supplementary Figure S13 for "PUMILIOs and m^6^A-ECT2/ECT3 share mRNA targets and exert opposing control over organogenesis"

**A**

| Accession | Gene | logFC | Padj | Description | |
|---|---|---|---|---|---|
| AT3G13470.1 | CPN60B2 | 4.010 | 2.03E-12 | TCP-1/cpn60 chaperonin family protein | Chaperone |
| AT2G28000.1 | CPN60A1 | 3.127 | 2.03E-12 | chaperonin-60alpha | |
| AT3G13860.1 | HSP60-3A | 3.040 | 2.03E-12 | heat shock protein 60-3A | |
| AT3G23990.1 | CPN60 | 2.224 | 3.98E-11 | heat shock protein 60 | |
| AT2G33210.1 | HSP60-2 | 1.375 | 2.77E-09 | heat shock protein 60-2 | |
| AT5G20720.1 | CPN20 | 1.066 | 1.78E-08 | chaperonin 20 | |
| AT1G67090.1 | RBCS-1A | 0.694 | 3.13E-06 | ribulose bisphosphate carboxylase small chain 1A | Photosynthesis |
| AT5G38410.3 | RBCS-3B | 0.621 | 2.40E-06 | ribulose bisphosphate carboxylase family protein | |
| ATCG00490.1 | rbcL | 1.110 | 4.56E-08 | ribulose-bisphosphate carboxylase | |
| AT1G12310.1 | CML13 | 0.828 | 1.69E-07 | Calcium-binding EF-hand family protein | $Ca^{2+}$ response |
| AT5G23060.1 | CAS | 0.816 | 7.40E-07 | calcium sensing receptor | |
| AT5G15200.1 | RPS9B | 0.816 | 1.55E-07 | Ribosomal protein S4 | Ribosome |

**B**

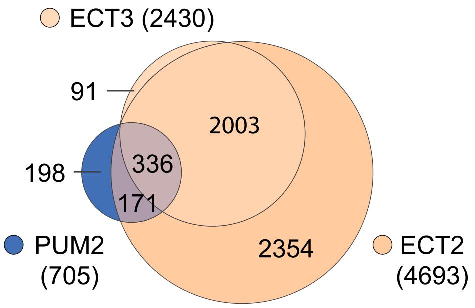


**C**

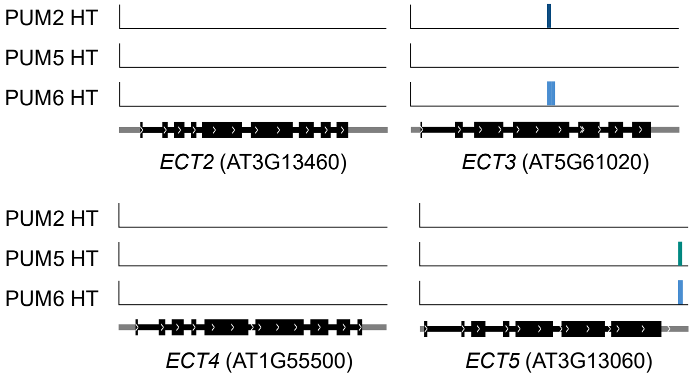


**Supplementary Figure S13. Lack of evidence for functional physical association between PUM and ECT proteins (extended data).**
**(A)** Identity of the 12 proteins enriched in both PUM2 IP-MS and ECT2 IP-MS (Tankmar et al., 2023). **(B)** Venn diagram showing overlaps in mRNA targets of ECT2, ECT3 and PUM2 identified by HyperTRIBE in aerial tissues. Chi-square test for mutual independence, $p < 2.2 \times 10^{-16}$. **(C)** Location of PUM2/5/6 HyperTRIBE editing sites on *ECT2/3/4/5* transcripts.
